# Inference of deformation mechanisms and constitutive response of soft material surrogates of biological tissue by full-field characterization and data-driven variational system identification

**DOI:** 10.1101/2020.10.13.337964

**Authors:** Z. Wang, J.B. Estrada, E.M. Arruda, K. Garikipati

## Abstract

We present a novel, fully three-dimensional approach to soft material characterization and constitutive modeling with relevance to soft biological tissue. Our approach leverages recent advances in experimental techniques and data-driven computation. The experimental component of this approach involves *in situ* mechanical loading in a magnetic field (using MRI), yielding the entire deformation tensor field throughout the specimen regardless of the possible irregularities in its three-dimensional shape. Characterization can therefore be accomplished with data at a reduced number of deformation states. We refer to this experimental technique as MR-***u***. Its combination with powerful approaches to inverse modelling, specifically methods of model inference, would open the door to insightful mechanical characterization for soft materials. In recent computational advances that answer this need, we have developed new, data-driven inverse techniques to infer the model that best explains the physics governing observed phenomena from a spectrum of admissible ones, while maintaining parsimony of representation. This approach is referred to as Variational System Identification (VSI). In this communication, we apply the MR–***u*** approach to characterize soft polymers regarding them as surrogates of soft biological tissue, and using VSI, we infer the physically best-suited and parsimonious mathematical models of their mechanical response. We demonstrate the performance of our methods in the face of noisy data with physical constraints that challenge the identification of mathematical models, while attaining high accuracy in the predicted response of the inferred models.

## 1 Introduction

Experimental measurements of full-field deformation are commonly restricted to in-plane or surface assessment of motion, e.g. with tracked deformation of a grid or speckle pattern [12, 7, 6, 34]. In many geometries, this can be considered representative of the through-thickness deformation (i.e. if plane stress or strain can be assumed) and a material sample can be inversely characterized accordingly. However, the plane stress or strain assumptions frequently break down in cases of biological materials, in which there are not only geometric complexities, but internal structural hierarchy and finite deformations as well. Thus, for applications aiming to characterize tissues, a fully three-dimensional measurement capability is a necessity. Furthermore, the class of three-dimensional imaging technique is an important consideration, as even if opacity is mitigated, conventional optical setups typically have a higher resolution in-plane versus out-of-plane. For these reasons, we have developed a magnetic resonance imaging- (MRI) based approach rooted in displacement-encoded MRI, in which alternating pulse field gradients in conjunction with synchronous sample motion and stimulated echoes provide phase-based images of the 3D displacement field [21]. This technique, termed alternating pulse field gradient stimulated echo imaging (APGSTEi), in a nod to the non-imaging variant pioneered by Cotts et al. [13], is presented in detail in Scheven et al. [32].

Given that testing of our class of samples necessitates measurement of full-field volumetric deformation for any combination of optical, mechanical, and/or biological material properties and geometry, we turn our attention to methodologies in which deformation measurements are translated into characterization directly. In general, this means solving the inverse problem—determining material properties by minimizing the error between the model solution and data, while respecting consistency with governing equations. One approach is to directly minimize the mismatch in the weak form of the momentum balance equation, the principle of virtual work. This class of material property identification techniques includes the constitutive equation gap method (CEGM) [15] and the virtual fields method (VFM) [26], which have been broadly expanded to a variety of applications in the past few decades including characterization of linear [24, 4, 16, 25, 40] and non-linear [17, 10] anisotropic metals and composites, hyperelastic rubbers [3, 27], and biological tissues [5, 30, 8, 41, 20]. In particular, the VFM assumes a constitutive form for the material model and iteratively adjusts material parameters within that constitutive law until the mismatch between the internal virtual work and applied external virtual work is minimized. Thus, for simple materials where the constitutive model is known, the virtual fields method is an excellent candidate for inverse characterization; the aforementioned APGSTEi technique is used as the imaging platform for VFM-based material characterization of silicone rubber samples in our prior work [14], and therein termed MR-***u***. However, soft and bio-materials have a variety of complicated material behavior and physics owing to chemistry and structural hierarchy; in many cases, the choice of material model is not firmly established. Furthermore, systematic and random error sources that require data filtering may in general have non-trivial implications in the resulting material parameter fits. In these cases – those where the underlying constitutive behavior and exact noise calibrations are not known *a priori* – the VFM is procedurally limited.

If our goal is not just to find parameter fits for a single chosen model and experimental noise correction method, but to describe the underlying physics with fidelity, we require a method that adjusts procedurally to reflect the true constitutive behavior itself. Hence, we expect to gain insight to the fundamental mechanisms by which native and engineered tissue undergo volumetric, isochoric, shear, isotropic and anisotropic deformation. These in turn, are critical for advances in tissue engineering, and orthopedic surgery. Therefore, we will adopt a novel method that can infer the proper physics from a large range of possible mechanisms, which is described next. This is the central thrust of the computational component of this work.

We have developed a class of inverse modeling techniques that allows the identification of physics from experimental data. Our first step in this direction was motivated by an interest in inference of mechanisms that govern the physics of dynamic pattern formation in materials. As background, these processes lead to microstructures via reaction-diffusion and phase transition phenomena. Our approach is based on the insight that, in any model, the forms of mathematical operators encode physical mechanisms. In the case of dynamic pattern formation in materials, the physics is governed by first-order (in time) partial differential equations–a class that includes the time-dependent reaction-diffusion and phase field equations. In this context, algebraic operators acting on concentrations represent (coupled) reactions between components of the material, while the differential, Laplacian operator represents Fickian diffusion. We have extended and exploited connections of other mathematical operators, including higher-order ones, to more complex physical mechanisms. The products of algebraic and differential operators reveal non-linearities, often introduced due to coupling that originates in the thermodynamics. Our aim is to infer the physics governing the observed phenomena by identifying the minimal set of operators whose combination into partial differential equations completely describes the data at hand. Our method works through the weak form of the governing system of equations. We therefore refer to this method as Variational System Identification, or VSI [37, 36, 38]. VSI uses step-wise regression, a statistical test, penalization on loss functions to control over-fitting, mathematically rigorous operator staggering and physically well-founded approaches to mechanism elimination.

VSI works with spatially well-resolved as well as sparse data that may be noisy. In more recent developments, we have extended VSI to problems in which the data are spatially non-overlapping at distinct times, and also are temporally sparse, as is the case with many techniques of materials processing and characterization. This extension leverages the definition of experimentally realizable quantities of interest by manipulation of the weak form, and, via consistency tests, provides mathematical guarantees on the inference of mechanisms [36]. A crucial ingredient in VSI is the recognition of universal aspects of the physics to intelligently constrain the paring down of operators. Thus, we have exploited the understanding of steady state *versus* transient phenomena [36], and here we extend it to physical stability of the inferred model. In related research, we also have developed Bayesian approaches to quantify the uncertainty in the inferred physics [38], and techniques that account for spatial and temporal heterogeneity of the physical mechanisms [39]. In this communication, we apply VSI to infer, from a range of possible candidates, the mechanisms of deformation of soft polymeric materials from MR-***u*** data.

Our approach to system identification is related to the Sparse Identification of Nonlinear Dynamics (SINDy) [9, 22, 31, 19], but differs in our use of the weak form rather than the strong form for identification of governing equations. VSI inherits many advantages from the weak form, including (a) the natural identification of boundary conditions, (b) introduction of basis functions to interpolate the data, which can be chosen to have high regularity, (c) transfer of derivatives to variations, thus lowering the regularity required in the data, and (d) isolating operators on the data by judicious choice of variations. A different approach to physics inference by Raissi and co-workers has focused upon determination of unknown coefficients in known systems of equations using Gaussian Processes [28] and deep neural networks [29]. However, in assuming the governing equations to have certain specific, classical forms, such as the Navier-Stokes equations, and seeking the corresponding coefficients, this approach takes a different path from ours. We seek to infer physics encapsulated in mathematical forms of equations and response functions: our methods, apart from obeying broad physical principles of conservation laws and symmetry, and exploiting transient *versus* steady state phenomena and stability, do not impose specific physics. Instead, we aim to infer the physical mechanisms of deformation by considering a wide range of them that are encompassed within the conservation laws and symmetry restrictions. Here, we extend our techniques for application to soft material surrogates for biological tissue. Finally, we note recent work in discovering governing equations in a constructive manner by combining graph theory with data-driven physics discovery [2]. An earlier approach of relevance in this spirit can be seen in Ref. [33].

The experimental methods are described in Section 2. Section 3 details the VSI technique. Our results for inferring deformation mechanisms of polymeric materials, as surrogates for soft biological tissue, from synthetic and experimental data appear in Section 4. A discussion and conclusions are presented in Section 5.

## 2 Methods

### 2.1 Full-field displacement capture (FFDC)

The acquisition of fully three-dimensional displacement fields from samples undergoing finite deformations was demonstrated by a combination of a custom magnetic resonance pulse sequence, cyclic actuated loading, and a subsequent data analysis procedure. The specific details of the apparatus and pulse sequence can be found in [32], while exact data processing procedures are presented in more detail in [14]. However, a brief procedure is reproduced for the ease of the reader below.

Material samples rigidly fixed on either end by two grips or platens are loaded in global uniaxial tension inside the center of a 7T small-animal MRI system. One end of the sample is fixed rigidly to the stationary end of a custom polyetherimide chamber, while the other side is permitted to move in the free axis by a prescribed global displacement. This displacement is set by a captive linear actuator (L5918S2008-T10X2-A50, Nanotec Electronic GmbH and Co. KG, Germany) acting at a sufficient distance from the MRI to avoid interaction with (and interference from) the strong magnetic field.

Samples are then loaded and imaged synchronously and cyclically. Global uniaxial deformation is applied to the sample within the linear range of the coil, which then is put into a state of tension and, at high loads, concurrently measured with a load cell (LCM300, Futek Advanced Sensor Technology Inc., Irvine, CA). For low forces (*<*10 N), loads at the equivalent global deformation states were measured externally and quasistatically using an ADMET materials testing system platform (ADMET, Inc., Norwood, MA, USA) equipped with a 25 N load cell. Briefly described, the imaging procedure proceeds by encoding magnetic spins into the sample in the deformed state via a set of alternating pulsed field gradients, storage of those spins during a macroscopic return of the sample to the reference configuration, and subsequently unencoding via a second, identical pulsed field gradient pair. The result is a 3D map of accumulated phase of voxels in the reference configuration during the course of a deformation cycle, corresponding to the chosen Cartesian encoding direction, and is wrapped to 2*π*. Repeating this process—encoding for each of the three Cartesian directions—with a linear conversion of phase to displacement results in the wrapped displacement field as a function of 3D position in the tested sample.

The stored phases corresponding to displacement can then be either unwrapped, e.g. via custom implementation of algorithms such as described in Ref. [1] or numerically differentiated directly by a division analogue to filter convolution (termed *divolution*), described in more detail in Ref. [14]. For the purposes of this study, the displacement fields are numerically differentiated and cast in the form of the deformation gradient tensor, from which other finite strain tensors may be determined in a straightforward manner (see Section 3 for details on the mathematical formulation). The choice of spatial filter on the displacement or deformation gradient data and when to apply it is, in general, left to the user; both thermal and systematic detector noise are practical considerations for the inclusion of specific filter types. Magnetic resonance, by virtue of being data acquisition in Fourier space, is particularly susceptible to Gibbs ringing artifacts, which are commonly improved by apodization, or using Hamming-type window filters in k-space. Thermal noise is another, distinct source of systematic experimental error which is improved by Gaussian filtering. The strengths of these filters in our prior study [14] and more general image processing applications commonly requires making a trade-off of sharpness and fidelity; light filtering can pick up sharper gradients at the expense of random noise while stronger filtering suppresses random noise while blurring boundary values. The exact filter sizes for an experiment are commonly left to the finesse of the user, but could in general be another variable over which we may optimize.

## 3 Inference of deformation mechanisms in soft materials by Variational System Identification

We describe VSI for the inference of constitutive laws from three-dimensional, full-field deformation data in soft materials obtained by the FFDC approach. Focusing on nonlinear hyperelasticity in this communication, we develop our methods in the context of the strain energy density function. In this setting, the relevant mechanisms of volumetric, isochoric, and anisotropic deformation are resolved by functions of the right Cauchy-Green strain tensor.

The displacement field is written as ***u***(***X***), where ***X*** is the reference position, the deformation gradient tensor is ***F*** = 1 + *∂u/∂****X***, and the right Cauchy-Green tensor is ***C*** = ***F*** ^T^***F***. It follows that volumetric response is entirely determined by *J* = det***F*** and the isochoric (volume preserving) response is determined by 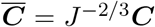. Defining the first principal invariant 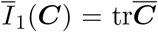, a commonly used constitutive model for isotropic and nearly incompressible soft biological tissue and polymers is the neo-Hookean strain energy density function,

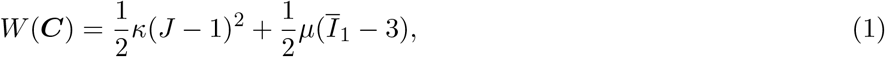

where the bulk modulus is *κ* and the shear modulus is *µ*. Our goal in this communication is to go beyond the limitation of such prescribed models. As prefaced in the Introduction, we now prepare to present VSI for the inference of models that represent the physics in sufficient detail (“expressiveness” in machine learning terms) while maintaining efficiency of representation. The building blocks of our approach are mathematical terms that encapsulate the relevant deformation mechanisms and lead to operators in the equations below.

VSI works by considering a number of deformation mechanisms as *possible* candidates in the strain energy density function. As candidate deformation mechanisms for soft biological tissue and polymers we consider volumetric, isochoric, and orthotropic mechanisms, via the additional principal tensor invariant 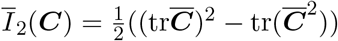, and 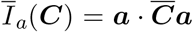 the squared stretch along a direction ***a***, by adding terms with coefficients *θ*_1_, *θ*_2_, … beyond those used in Equation (1):

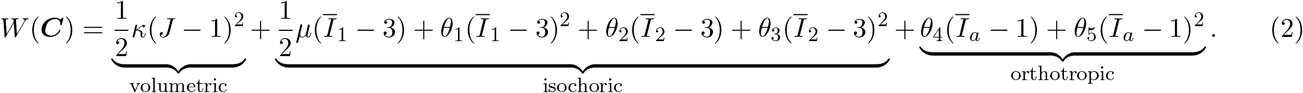

The first Piola-Kirchhoff stress tensor is ***P*** = *∂W/∂****F*** and leads to the form

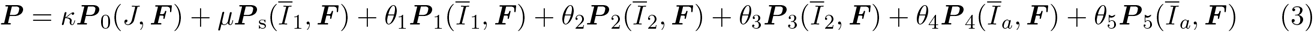

For the sake of brevity, we have omitted the explicit forms of the stress operators ***P***_0_–***P***_5_. However, they are straightforwardly recognized as the derivatives with respect to ***F*** of the functions with coefficients *κ, µ, θ*_1_, …, *θ*_5_ in Equation (2). VSI works with the weak form of the stress equilibrium equation, written as:

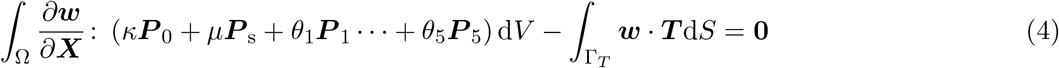

where ***w*** is the weighting function (the variation on the displacement field) and we have used Equation (3) for the stress. The solid volume is represented by Ω and Γ_*T*_ is the part of the boundary over which traction loading, ***T***, is applied.

Given FFDC data as a set of three-dimensional displacement vectors, ***d***_*i*_, where *i* = 1, …, *N* enumerate the points ***X***_*i*_ throughout the reference configuration of the sample where data has been obtained, we construct a finite element mesh with nodes at ***X***_*i*_. Using appropriate finite element interpolation functions, *N* ^*A*^(***X***), where *A* = 1, …, *n*_nel_ runs over the number of nodes in the element, we construct the displacement field ***u*** and weighting function, ***w***, defined within each element (subscripts/superscripts denoting elements are suppressed),

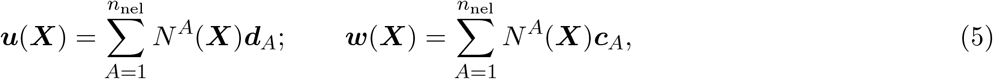

where, as is well-known in finite element methods, ***c***_*A*_ are arbitrary nodal weighting function values. For an element node *A* located at ***X***_*i*_, we have ***d***_*A*_ = ***d***_*i*_. The element deformation gradient tensor ***F***, and the kinematic quantities *J*, 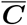, *Ī*_1_, *Ī*_2_ and *Ī*_*a*_ can all be computed from the data using the finite element interpolations in Equation (5).

Using Equation (5) in Equation (4) we arrive at the Galerkin weak form in terms of the stress operators, now written as 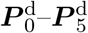 with superscripts (*•*)^d^ emphasizing that the stress is computed from the known *data*.

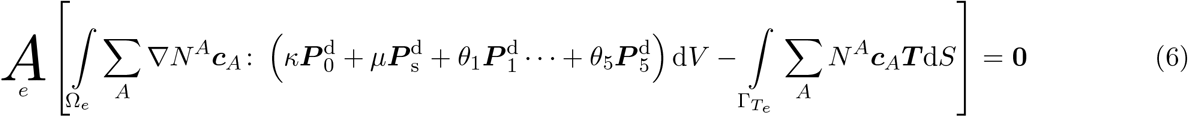

where 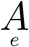 is the finite element assembly operator over elements. Additionally, 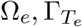 denote an element sub-domain and traction boundary restricted to an element, respectively.

Following the standard procedure in finite element computations, Equation (6) leads to the residual vector after accounting for the arbitrariness of weighting function degrees of freedom:

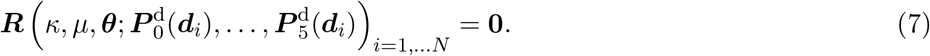

The above notation indicates that ***R*** has a *functional dependence* on *κ, µ*, and the additional coefficients ***θ*** = {*θ*_1_, … *θ*_5_}. Also note that since the stress is *parameterized* by the data ***d***_*i*_ via the operators 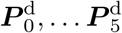, we have chosen to highlight this dependence directly, rather than through the deformation gradient and other kinematic quantities as in Equation (3). The residual vector is assembled from the element contributions and is defined here as

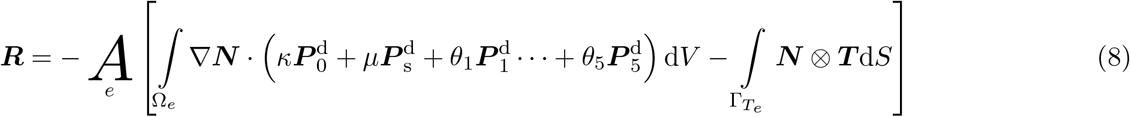

where 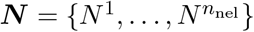 is the vector of interpolation functions in an element.

### 3.1 Basis operators

VSI associates deformation mechanisms with basis operators that are written in weak form. The known boundary traction term serves as the label in machine learning parlance, and written as ***y***:

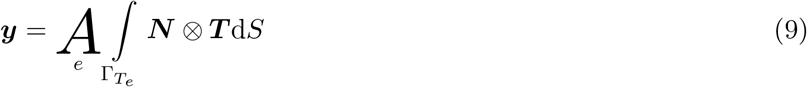

The Piola-Kirchhoff stress contributions arising from the various terms in the strain energy density lead to matrix operators **Ξ**_*α*_ that we express in the general form:

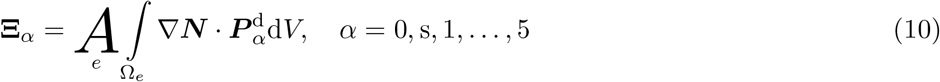

The residual ***R*** then follows from Equations (9) and (10) as

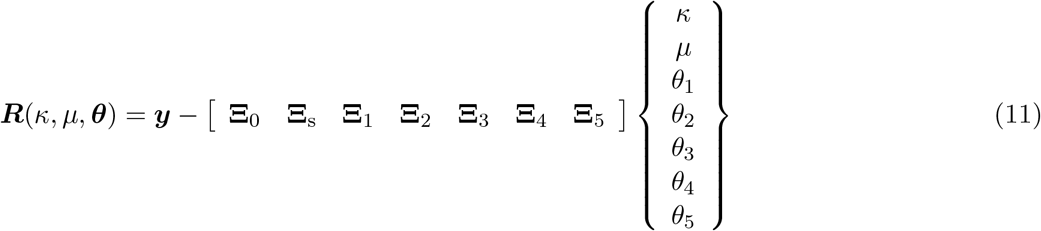

We write ***θ*** = ⟨*θ*_1_, …, *θ*_5_ ⟩^T^, but in the interest of transparency, we continue to delineate the bulk and shear moduli, *κ* and *µ* in what follows. We also use **Ξ** = [**Ξ**_0_ **Ξ**_s_ **Ξ**_1_ **Ξ**_2_ **Ξ**_3_ **Ξ**_4_ **Ξ**_5_].

VSI enables physics inference by eliminating some and accepting other deformation mechanisms, and thus reducing the coefficients *κ, µ*, ***θ*** to a minimal set of non-zero values. A loss function, 𝓁 is defined as

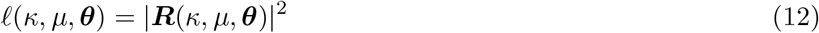

Ridge regression is applied with regularization of the coefficients via a penalty term to prevent overfitting

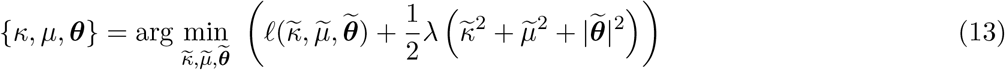

where *λ* is the penalty coefficient. This step yields the coefficient vector in the form:

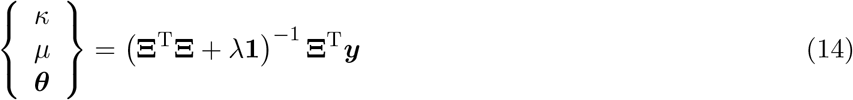

### 3.2 Stepwise regression

This approach consists of an iterative procedure that is combined with standard linear regression to identify and eliminate inactive deformation mechanisms by setting their coefficients to vanish in the set {*κ, µ*, ***θ***}. The corresponding algorithm for standard regression uses the statistical *F* -test (explained below and presented in schematic form in Figure 2), which evaluates the significance of the increase in loss, 𝓁 (*κ, µ*, ***θ***) relative to the gain in efficiency of representation [37]. This approach is formally equivalent to using sparsity-inducing norms on the loss function, such as the *L*^1^-norm coupled with cross-validation or LASSO (least average shrinkage and selection operator; a statistical and machine learning algorithm that performs regression while combining variable selection and regularization). However, we have found step-wise regression to perform better, and the use of the other methods to need insightful choices of priors for the coefficients (*κ, µ*, ***θ***). We use backward model selection by stepwise regression [18], a strategy of starting with a large library of models and culling them. As we have demonstrated previously, this approach delivers parsimonious results with VSI [37, 36]. The algorithm is summarized below.

#### Algorithm 1: Stepwise regression with the *F* -test

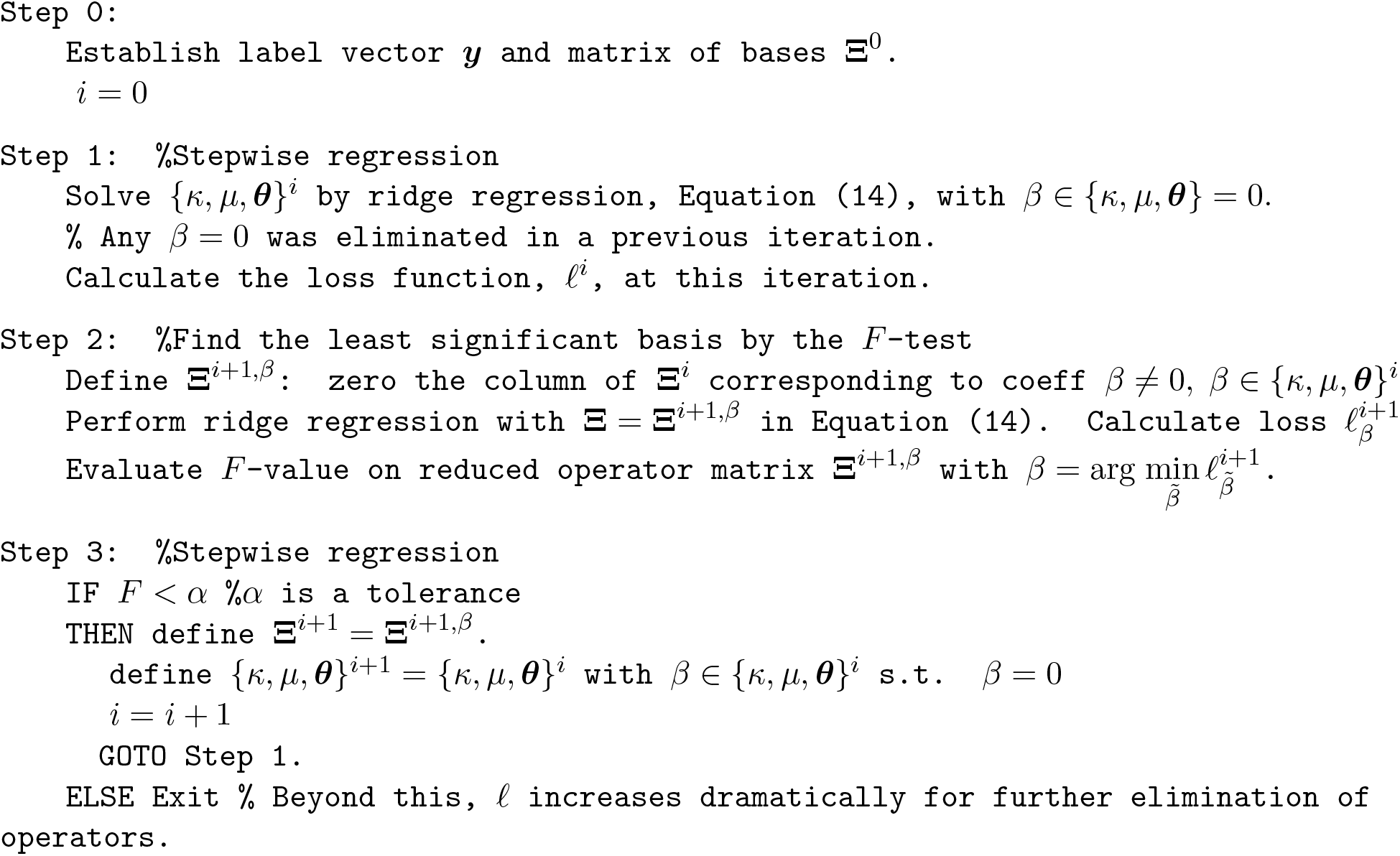

**Figure 1:**
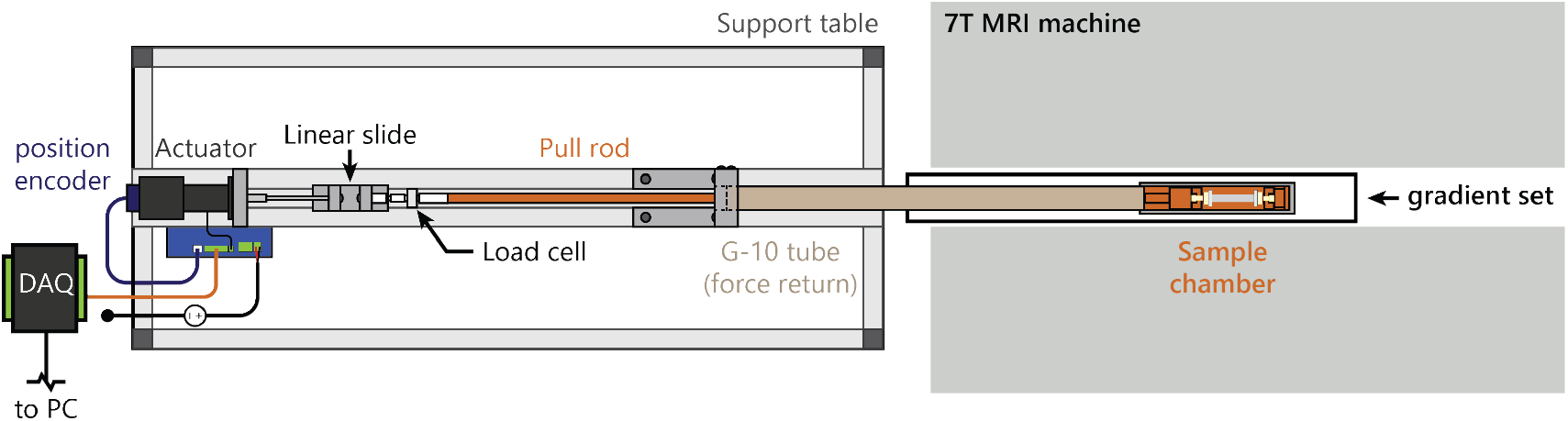
Schematic of the experimental setup (top view) for full-field displacement capture. A linear actuator prescribes a periodic, constant-amplitude, encoder-verified displacement of one edge of a sample, synchronous with a custom nuclear magnetic resonance pulse sequence which acquires slices of the entire full-field 3D displacement between reference states. A load cell measures the applied force on the sample, which is balanced by a force-return G10 tube and custom polyetherimide (Ultem) sample chamber [14].

**Figure 2:**
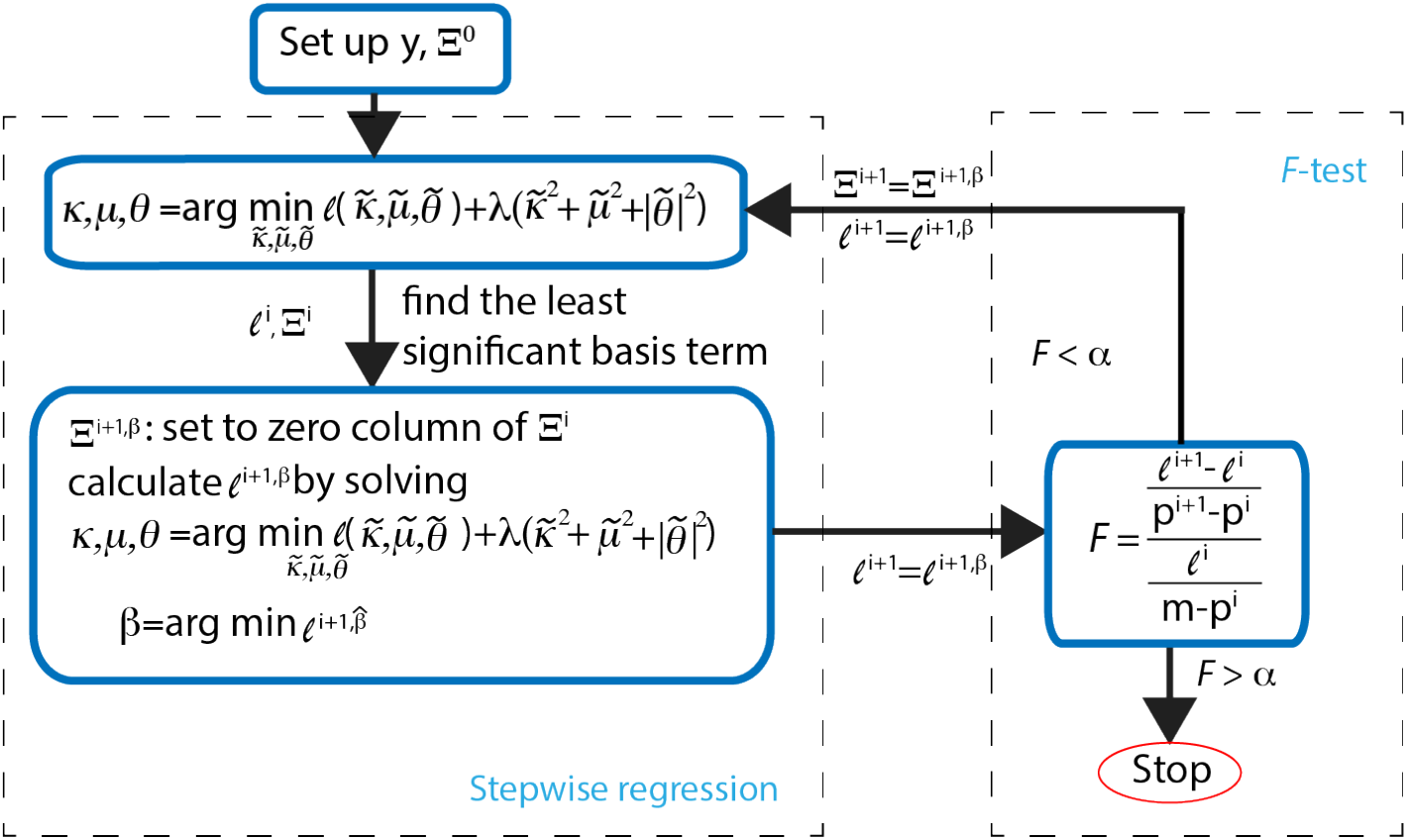
Schematic of Algorithm 1 for stepwise regression with the *F* -test.

There are several choices of the criterion for eliminating deformation mechanisms via the operators in **Ξ**. Here, we adopt a widely used statistical criterion called the *F* -test, also used by us previously [37, 36]. The significance of the change between the model at iterations *i* + 1 and *i* is evaluated by:

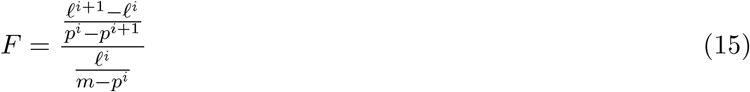

where *p*^*i*^ is the number of operators at iteration *i* and *m* = 7 is the total number of operators. The *F* test normalizes the increase in loss function by the gain in efficiency of representation (decreasing model complexity). If *F* exceeds a pre-defined tolerance, *α* (Step 3 in Algorithm 1), stepwise regression terminates without further elimination of operators.

The penalty coefficient in ridge regression,^1^ *λ*, is chosen to lie in the range [1, 10] by leave-one-out crossvalidation at each iteration.^2^ We illustrate Algorithm 1 for stepwise regression algorithm with the *F* -test in Figure 2.

Some remarks are relevant on the use of the weak form in VSI. As is well-known, the weak form transfers spatial derivatives from the differential operators acting on the data to the weighting function in Equation (4). Combined with the use of smooth basis functions (such as splines) in Equation (5), and in the weak form Equations (6-7), this contributes to the robustness of VSI when confronted by noisy data [37, 36], relative to other related techniques that use the partial differential equation in strong form [9, 31, 11]. As we have shown in Ref. [37], VSI also is able to identify boundary conditions on the data, which the other techniques cannot.

### 3.3 Staggered VSI with physically based operator suppression

Noise in the FFDC data can mask physically meaningful constraints, such as near-incompressibility of the response (*J ∼* 1). We have developed a staggered approach to inferring the deformation mechanisms in order to circumvent this difficulty: In a first step we exclude the volumetric term from the mechanisms in Equation (2), effectively assuming *J* = 1. With the traction loading term as the label, we collect into ***y***^surf^ only those elements of the residual vector that correspond to boundary degrees of freedom, noting that DIM[***y***^surf^] *<* DIM[***R***]. We gather the rows of **Ξ**_s_, **Ξ**_1_, …, **Ξ**_5_ that correspond to boundary degrees of freedom into **Ξ**^surf^ and the others in its complement **Ξ**^in^, and similarly partition ***R*** into ***R***^surf^ and ***R***^in^ to rewrite Equation (11) as

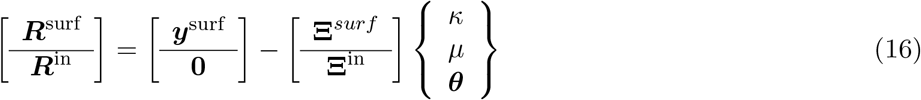

For the reduced vector of coefficients with ⟨0, *µ*, ***θ*** ⟩^T^, we first carry out VSI via Algorithm 1 with the residual ***R***^surf^ given by

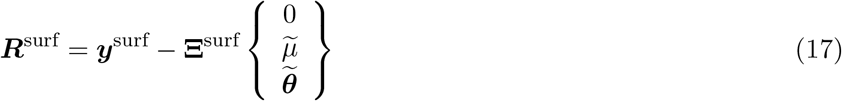

instead of ***R***. In this step we detect the active operator set consisting of **Ξ**^surf^, paired with predictor coefficient values 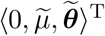. The volumetric operator is excluded at this step since the corresponding coefficient is set to zero.

We next reintroduce the volumetric operator with coefficient 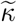 to the interior part of the residual, which is then reduced to:

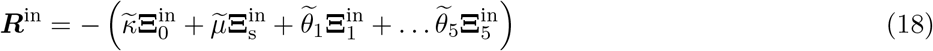

Holding 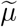 and 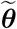 fixed, and using ridge regression, we find a predictor for 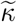. At this stage of the staggered VSI algorithm, we note that with the predictor coefficient set 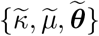 the surface residual 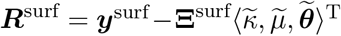 yields a value for |***R***^surf^| that exceeds the value attained by stepwise regression with ***R***^surf^ defined in Equation (17). We therefore consider a linear scaling, 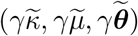, thus similarly scaling |***R***^in^| as defined by Equation (18), but not affecting its minimizer. However, it allows us to find a new minimum for |***R***^surf^| in the full coefficient space. Returning to the upper row block form of Equation (16), we impose 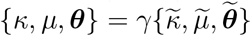 in

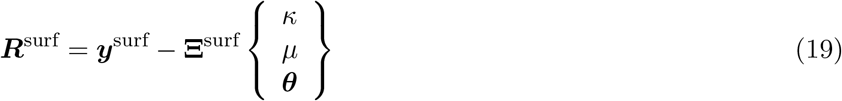

to find *γ* that thus defines a constrained minimizer *{κ, µ*, ***θ****}* of |***R***^surf^| by Algorithm 1. We thus attain minima of |***R***^surf^| and |***R***^in^|, and because of the quadratic dependence on (*κ, µ*, ***θ***), of |***R***| as well. This staggered approach to stepwise regression leads to robust VSI in the incompressible limit even with noisy data.

#### 3.3.1 Operator suppression

The available data may be restricted to a regime in which some of the operators arising from the deformation mechanisms in Equation (2) display near-linear dependence. The inferred set {*κ, µ*, ***θ***} can then harbor nonphysical results such as a negative bulk modulus, *κ*, or a vanishing shear modulus, *µ*. Recognizing *κ, µ* ≤ 0 as a loss of pointwise mechanical stability, this suggests an approach in which only one of the operators with linear dependence be admitted while suppressing the others in implementing the above staggered VSI. This represents the generally applicable principle of stability as a physics constraint incorporated within the inference techniques that constitute VSI. In a related approach, we have previously used the distinction between steady state and transient regimes of problems with first-order dynamics as a principle to strengthen VSI inference [36], and the stability-based criterion is in a similar spirit. We list the algorithm below:

##### Algorithm 2: Staggered VSI with operator suppression

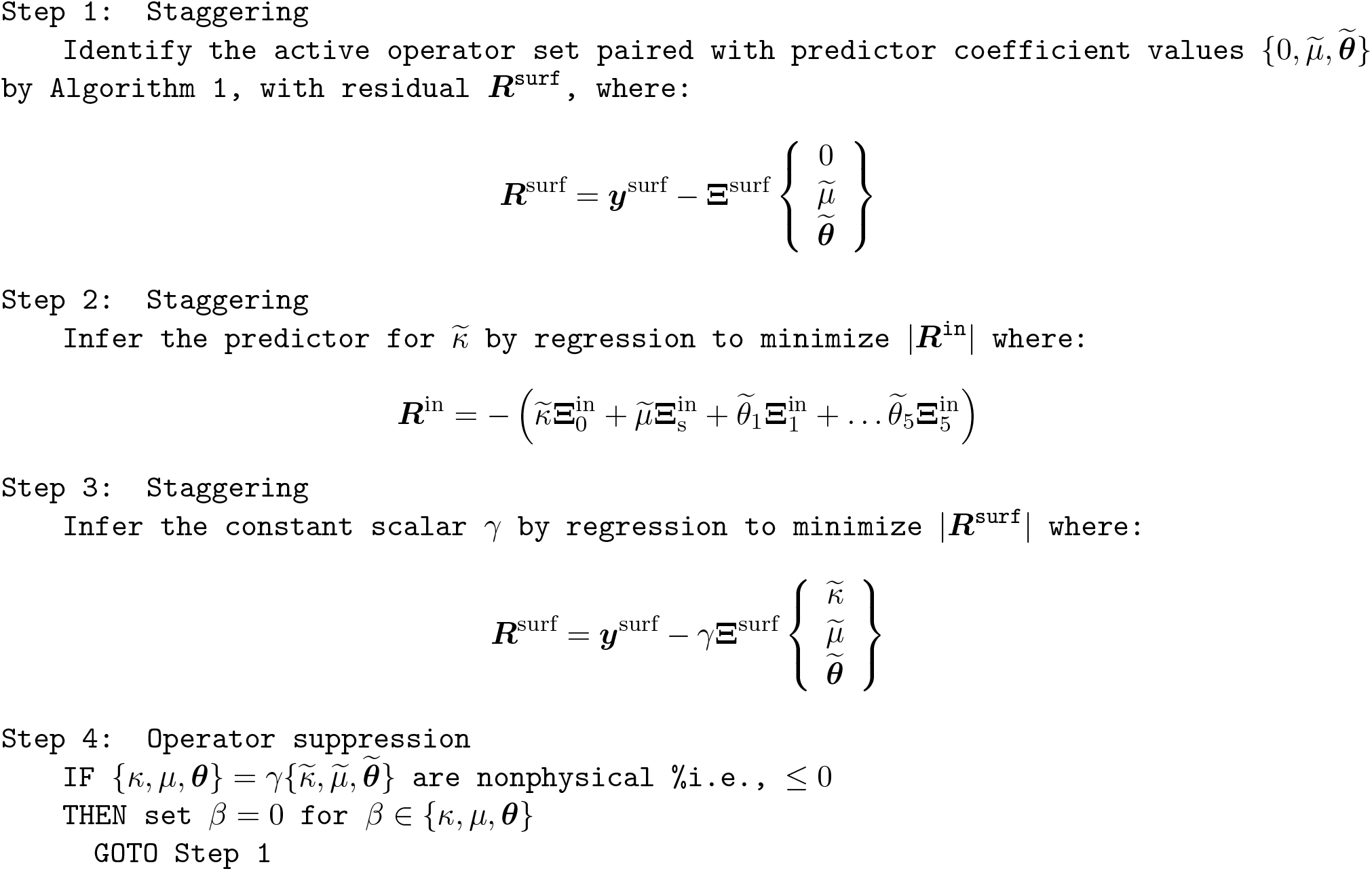

#### 3.3.2 Variational System Identification in comparison with the Virtual Fields Method

Variational System Identification and the Virtual Fields Method are both built on the weak form of the governing equation. Apart from this common foundation, however, the methods differ. VFM uses a number of virtual fields, or weighting functions in the terminology of the weak form, to explore an assumed constitutive model. Instead, VSI, following the standard approach in variational methods, extends the imposition of the weak form to its discretized version, thus arriving at a single equation that holds across weighting functions. At this step, VSI includes a library of candidate operators from which the model is to be inferred. VFM optimizes coefficients of the assumed model over all evaluations of the weak form for the applied virtual fields. In this sense, it bears similarities with PDE-constrained optimization methods, which however, may use different cost functions with additional regularization on the inferred coefficients, as well as a range of optimization algorithms. In contrast, given the discretized residual vector, Equation (8), VSI seeks to iteratively find optimal coefficients by driving the norm of the residual to vanish, while additionally eliminating operators corresponding to the least significant mechanisms and the corresponding coefficients. By using stepwise regression for this purpose, VSI goes beyond optimization of an assumed model to cull contributions from the larger library of operators. This strategy for choosing operators or mechanisms from a larger set of candidates is a feature not common in traditional optimization approaches.

### 3.4 PDE-constrained minimization following VSI

After the deformation mechanisms are identified by Algorithm 1, with Algorithm 2 applied as needed, the preliminary, predicted values of coefficients corresponding to the identified operators can be further refined by minimizing the error on ***u*** subject to the identified physics:

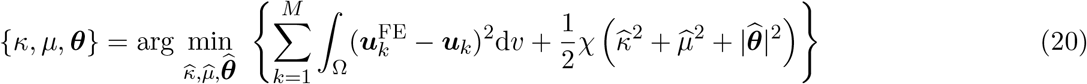

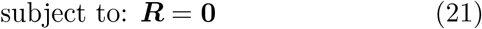

where *M* is the total number of loading cases, 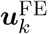 is the displacement field from the forward finite element solution obtained with the current values of coefficients 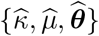 for the *k*th load step, ***u***_*k*_ is the corresponding FFDC data and *χ* is a penalty coefficient for regularization. Such PDE-constrained optimization needs the gradient of the corresponding loss function, 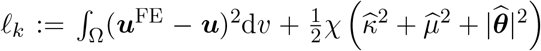 with respect to the parameters 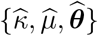. One approach to approximating the gradient is by finite differences, which requires a large number of solutions of the PDE constraint to construct numerical derivatives. Here, we adopt the adjoint method to evaluate the gradient, which requires a single solve of the (linear) adjoint equation of the original PDE constraint and a nonlinear solution for the parameter set 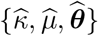 .

Since this is a standard implementation of adjoint-based gradient optimization with PDE constraints, we do not elaborate on it any further. In this work we use the L-BFGS-B optimization algorithm from the SciPy package [35] and the dolfin-adjoint software library [23] to solve the PDE-constrained minimization problem.

## 4 Results

### 4.1 Synthetic data

We first applied VSI to synthetic data generated from forward computations on rectangular prismatic blocks (42 × 6 × 6 mm^3^). We used the neo-Hookean strain energy density function, Equation (1), with true values of the coefficients appearing in Table 1. Note that with these coefficient ranges the neo-Hookean model represents a spectrum of soft biological tissue and polymers. Also included is the Poisson ratio that would be computed for the linear regime of infinitesimal strain with this model. As this initial Poisson ratio increases from *ν* = 0.495 to *ν* = 0.4995, the response tends toward near-incompressibility. We generated synthetic data for four different deformation modes: (a) Extension 1, loaded by a uniformly distributed tensile traction on the face with outward normal ***e***_1_ and fully fixed on the face with outward normal −***e***_1_. (b) Extension 2, loaded by a uniformly distributed tensile traction on the face with outward normal ***e***_2_ and fully fixed on the face with outward normal −***e***_2_. (c) Bending, loaded by a uniformly distributed shear traction along ***e***_3_ on the face with outward normal ***e***_1_ and fully fixed on the face with outward normal −***e***_1_. (d) Torsion, loaded by a twisting moment on the face with outward normal ***e***_1_ and fully fixed on the face with outward normal −***e***_1_. These loads resulted in maximum displacements over the respective domains of ∼ 8, ∼ 3, ∼ 10 and ∼ 4 mm. Forward solutions were obtained for these four boundary value problems using a three-field, Hu-Washizu finite element formulation in order to treat the response of the nearly-incompressible materials. The deformed configurations appear in Figure 3. We collected these displacement solutions, and then added noise with magnitude drawn from a Gaussian distribution with zero mean and standard deviations *σ* = 10^−3^ mm and *σ* = 10^−4^ mm, consistent with a thermal noise profile in displacement-encoded MRI. We thus have one noise-free data set and two noisy data sets for each case, leading to 24 data sets in total for the two different materials from four different boundary value problems. Note that the relative errors of the corrupted displacement are larger near locations with smaller boundary values. However the proposed bases, characterized by three invariants of ***C***, which are effectively determined by the gradient of displacement, may not be small at these locations. In order to correct for the artifact of large amplification of noise relative to the label, we have not considered spatial points where any of the candidate operators evaluated below a threshold, *ϵ*.

**Table 1:**
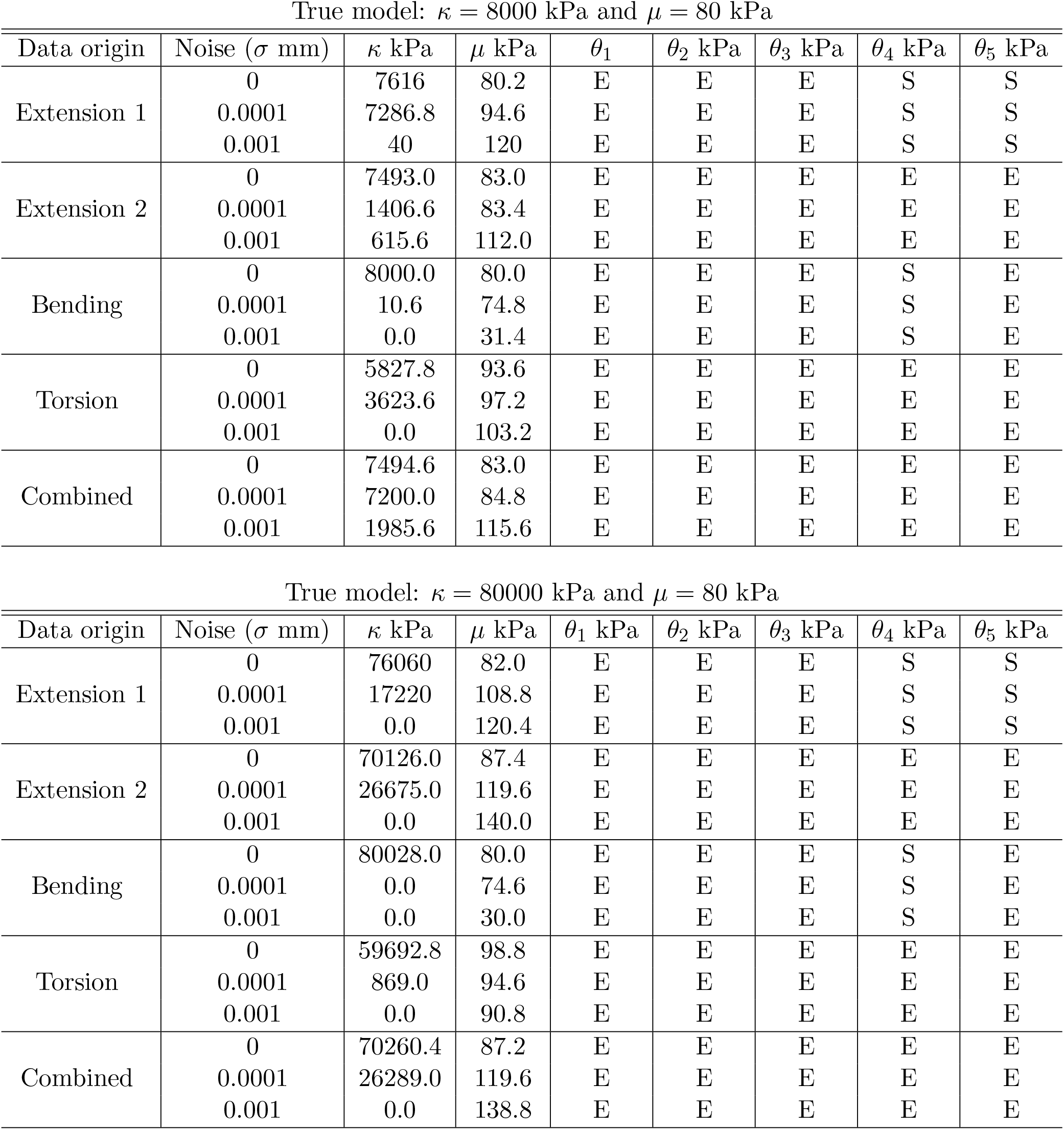
VSI results for synthetic data with bulk modulus *κ* = 8000 kPa, shear modulus *µ* = 80 kPa (Poisson ratio *ν* = 0.495 in the infinitesimal regime) and *κ* = 80000 kPa, shear modulus *µ* = 80 kPa (Poisson ratio *ν* = 0.4995 in the infinitesimal regime). These two cases test VSI for the progression toward incompressibility *ν* → 0.5.”E” denotes that the corresponding operator is eliminated by VSI, and “S” denotes the operator is suppressed to prevent a material instability.

**Figure 3:**
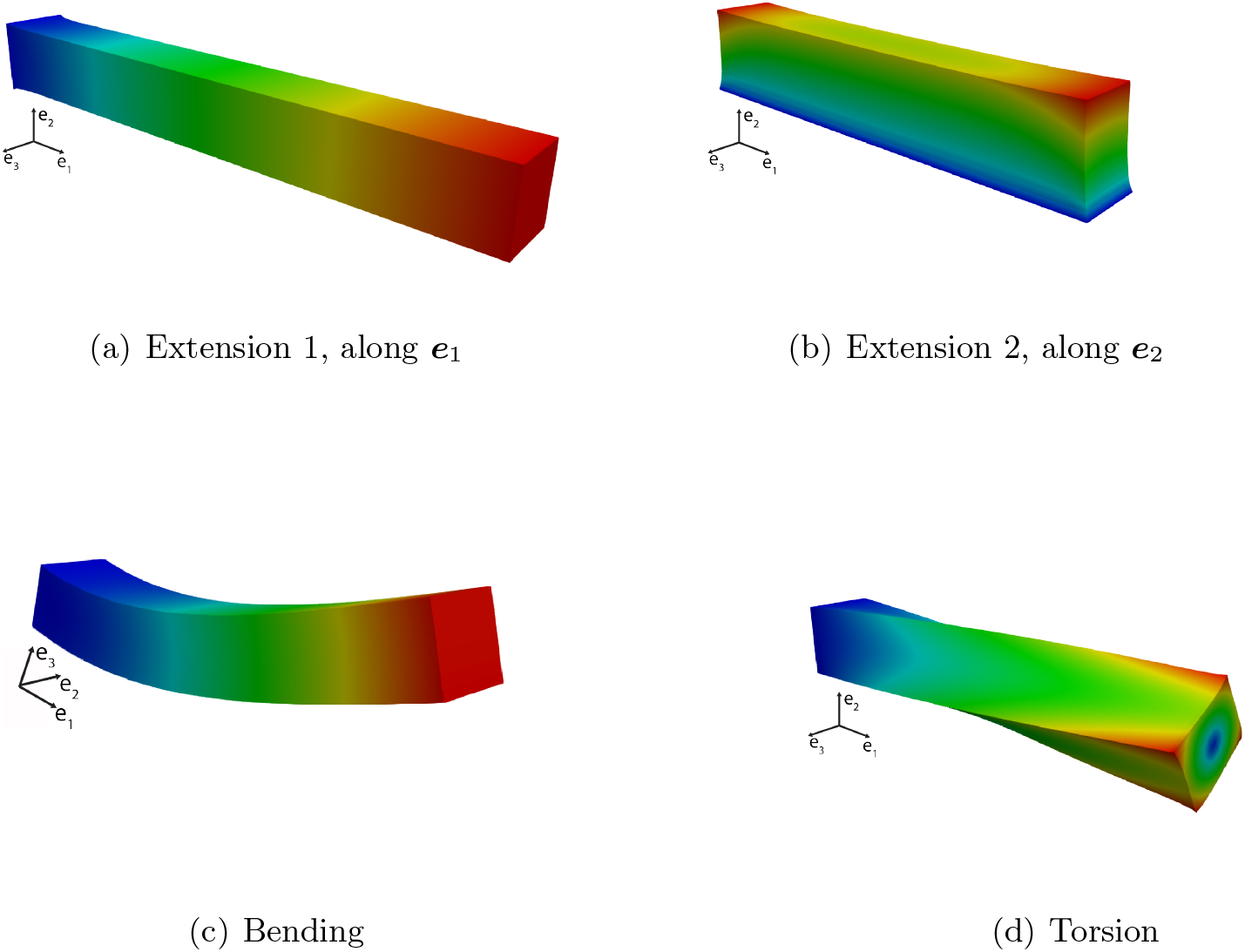
The four boundary value problems for generation of synthetic data.

VSI was applied starting with all the deformation mechanisms; i.e., without suppression of any of the operators, and with the anisoptropy direction ***a*** = ***e***_1_. Figure 4 illustrates the functioning of Algorithms 1 and 2 of VSI via a stem and leaf plot and the iterate-wise loss curve for the noise-free synthetic data. Step 1 of staggering (Algorithm 2) is shown, for which reason only the updated values of *µ, θ*_1_ … *θ*_5_, are shown at each iteration. On the right in Figure 4, the plot shows the iterate-wise loss remaining low until a single mechanism, corresponding to the shear coefficient *µ*, remains. Steps 2 and 3 of Algorithm 2 provide the final values for *κ*. Using data from Extension 1 and Bending, incorrect mechanisms are identified, as reflected by a vanishing shear modulus corresponding to a loss of material stability and a nonphysical model. This is because the two orthotropic terms, corresponding to (*Ī*_*a*_ − 1) and (*Ī*_*a*_ − 1)^1^ components in the candidate free energy density function (Equation 2), display near linear dependence with the shear term as shown in Figure 5. However, using Algorithm 2 for staggering and suppression of the anisotropic operators, VSI was successful in identifying a physically meaningful set of deformation mechanisms even in the incompressible limit with noise, although the loss in accuracy of coefficients increases with the approach to incompressibility and higher noise (Shown in Table 2). For Extension 2 and Torsion, the correct deformation mechanisms were identified without operator suppression due to the linear independence of the deformation mechanisms as shown in Figure 5. Notably, when all data were combined together and used, we were able to re-identify the correct mechanisms without operator suppression.

**Table 2:** Mechanism coefficients identified by VSI in Table 1 now refined by PDE-constrained minimization. The two values of bulk modulus *κ* = 8000 kPa, shear modulus *µ* = 80 kPa (Poisson ratio *ν* = 0.495 in the infinitesimal regime) and *κ* = 80000 kPa, shear modulus *µ* = 80 kPa (Poisson ratio *ν* = 0.4995 in the infinitesimal regime) demonstrate results for the progression toward incompressibility *ν* → 0.5

| True model: $W = 4000(J - 1)^2 + 40(\bar{I}_1 - 3)$ kJm$^{-3}$ | | |
| --- | --- | --- |
| Data origin | Noise ($\sigma$ mm) | Identified strain energy density function (kJm$^{-3}$) |
| Extension 1 | 0 | $W = 4000.00(J - 1)^2 + 40.00(\bar{I}_1 - 3)$ |
| | 0.0001 | $W = 3999.99(J - 1)^2 + 40.00(\bar{I}_1 - 3)$ |
| | 0.001 | $W = 4000.13(J - 1)^2 + 40.00(\bar{I}_1 - 3)$ |
| Extension 2 | 0 | $W = 4000.02(J - 1)^2 + 40.00(\bar{I}_1 - 3)$ |
| | 0.0001 | $W = 4000.02(J - 1)^2 + 40.00(\bar{I}_1 - 3)$ |
| | 0.001 | $W = 3998.80(J - 1)^2 + 40.00(\bar{I}_1 - 3)$ |
| Bending | 0 | $W = 4000.00(J - 1)^2 + 40.00(\bar{I}_1 - 3)$ |
| | 0.0001 | $W = 4099.36(J - 1)^2 + 39.46(\bar{I}_1 - 3)$ |
| | 0.001 | $W = 3999.99(J - 1)^2 + 40.00(\bar{I}_1 - 3)$ |
| Torsion | 0 | $W = 4000.00(J - 1)^2 + 40.00(\bar{I}_1 - 3)$ |
| | 0.0001 | $W = 3999.74(J - 1)^2 + 40.00(\bar{I}_1 - 3)$ |
| | 0.001 | $W = 3979.70(J - 1)^2 + 40.00(\bar{I}_1 - 3)$ |

| True model: $W = 40000(J - 1)^2 + 40(\bar{I}_1 - 3)$ kJm$^{-3}$ | | |
| --- | --- | --- |
| Data origin | Noise ($\sigma$ mm) | Identified strain energy density function (kJm$^{-3}$) |
| Extension 1 | 0 | $W = 39996.62(J - 1)^2 + 40.00(\bar{I}_1 - 3)$ |
| | 0.0001 | $W = 3999.49(J - 1)^2 + 40.00(\bar{I}_1 - 3)$ |
| | 0.001 | $W = 39991.62(J - 1)^2 + 40.00(\bar{I}_1 - 3)$ |
| Extension 2 | 0 | $W = 39924.27(J - 1)^2 + 40.01(\bar{I}_1 - 3)$ |
| | 0.0001 | $W = 39845.05(J - 1)^2 + 40.01(\bar{I}_1 - 3)$ |
| | 0.001 | $W = 39430.81(J - 1)^2 + 40.04(\bar{I}_1 - 3)$ |
| Bending | 0 | $W = 40000.00(J - 1)^2 + 40.00(\bar{I}_1 - 3)$ |
| | 0.0001 | $W = 39629.32(J - 1)^2 + 40.66(\bar{I}_1 - 3)$ |
| | 0.001 | $W = 39813.08(J - 1)^2 + 40.31(\bar{I}_1 - 3)$ |
| Torsion | 0 | $W = 39604.10(J - 1)^2 + 40.02(\bar{I}_1 - 3)$ |
| | 0.0001 | $W = 37966.09(J - 1)^2 + 40.10(\bar{I}_1 - 3)$ |
| | 0.001 | $W = 36507.23(J - 1)^2 + 40.17(\bar{I}_1 - 3)$ |

**Figure 4:**
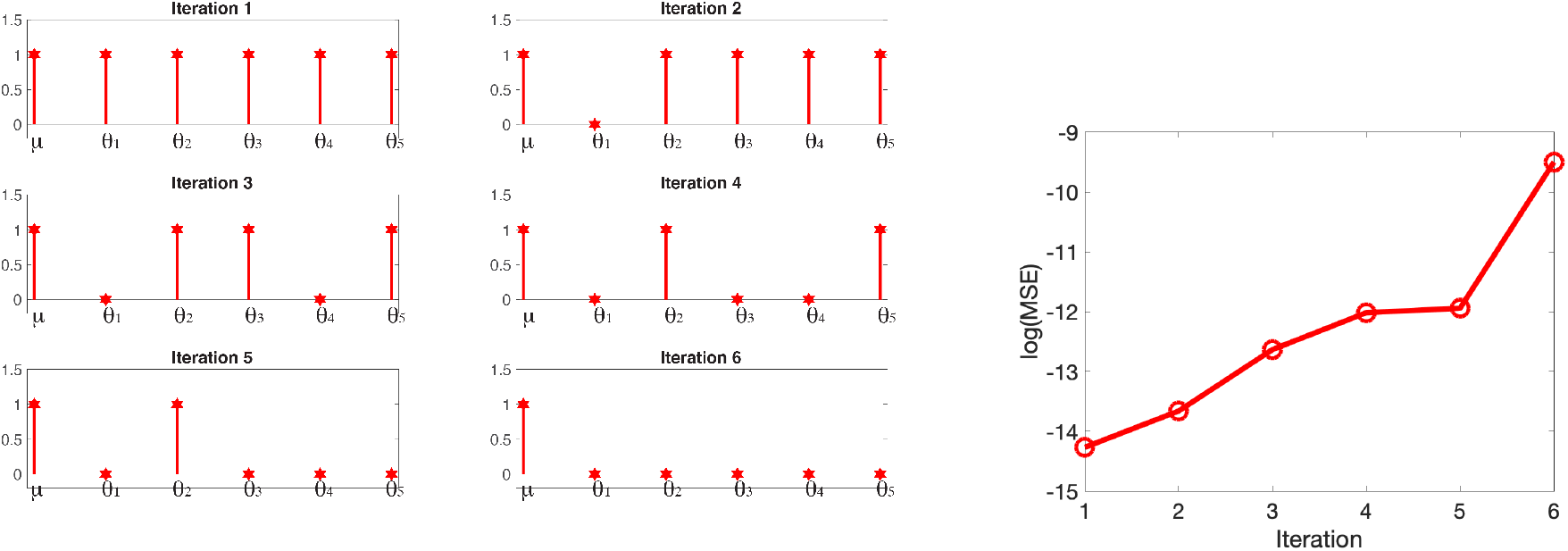
Left: Stem and leaf plot demonstrating step-wise regression via Algorithms 1 and 2 in staggered VSI: The coefficients *µ, θ*_1_, …, *θ*_5_ correspond to the non-volumetric terms in Equation (2). VSI sets coefficient *θ*_1_ to zero in Iteration 2, and the algorithm proceeds until *θ*_2_ is set to zero in Iteration 6. Right: the loss corresponding to each iteration of step-wise regression.

**Figure 5:**
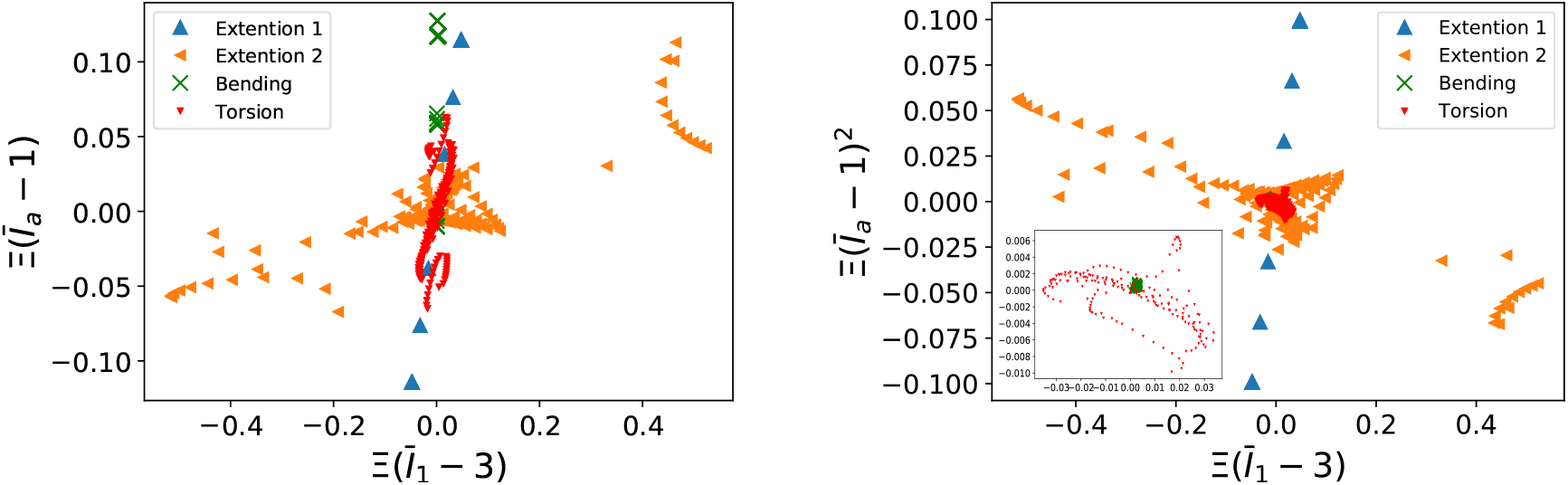
Synthetic data for the orthotropic terms *versus* the shear term in the candidate free energy density function evaluated for the four boundary value problems. The left plot shows that the orthotropic term, (*Ī*_*a*_ − 1), displays near linear dependence on, and even proportionality with, the basis term corresponding to the *Ī*_1_ − 3 component for Extension 1 and Bending. But the near linear dependence does not appear in Extension 2 and Torsion. The right plot shows that (*Ī*_*a*_ − 1)^2^ displays near linear dependence on *Ī*_1_ − 3 for Extension 1 but not for the other boundary value problems. The inset in the right plot is a magnified view of the Bending and Torsion data, which fall in very narrow ranges. Also note that for Extension 1, the data at each of six load steps nearly coincide for all reported spatial positions.

Due to the nearly-incompressible response of the these boundary value problems, *J* − 1 ∼ 0, and inference of the volumetric term can be significantly affected by noise. Imposition of non negative coefficients as a physical constraint can drive the bulk modulus to zero when using the data with higher amounts of noise. This is consistent with the finding in Ref. [14] that the estimated value for the bulk modulus may be nonphysically low or zero for near-incompressible materials. However, the staggered approach in Algorithm 2 guards against this pathology.

Following inference of the deformation mechanisms by VSI, the coefficient values, specifically the bulk and shear moduli *κ* and *µ*, were further refined by PDE-constrained minimization with adjoints as described in Section 3.4. We obtained coefficients with very high accuracy as shown in Table 2. VSI combined with PDE-constrained minimization thus leads the successful inference of the deformation mechanisms on synthetic data.

### 4.2 FFDC data from uniaxial loading on a soft elastomer

We next applied VSI to MR-***u*** data obtained from uniaxial loading on a soft elastomer as a surrogate for biological tissue. A rectangular prismatic block (41.5 × 8 × 7.5 mm^3^) of a commercially available elastomer (10A formulation; Dragon Skin, Smooth-On Inc., Macungie, PA) glued between two platens was subject to fully constrained displacement boundary conditions on one face, and stretched longitudinally in nine increments by specifying the maximum displacement (on the opposite face) to be 0.55, 1.09, 2.14, 3.25, 4.31, 4.89, 5.43, 6.46 and 7.57 mm. The resultant forces were: 0.16, 0.30, 0.58, 0.84, 1.09, 1.21, 1.32, 1.54 and 1.74 N, respectively, and had nearly uniform traction distribution for the constrained uniaxial loading. The details of the experimental configuration can be found in Ref. [14].

The reference configuration and reconstructed deformed configuration for the ninth load increment (of nine) appear in the left plot in Figure 6. A very slight softening response can be observed. Note that in keeping with convention, the results are reported as applied load *versus* the tip displacement. The FFDC data were obtained over a field of size 95 × 13 × 13 voxels. Field data were obtained for the displacement and for the deformation gradient tensor using the FFDC technique and smoothed by Hamming filters (which are typical for Gibbs ringing reduction in MR image processing) with different kernel sizes. As an alternate approach to smoothing, we fit the displacement data to polynomials and obtained a global quadratic field. The FFDC data as displacements and deformation gradient tensors, subject to these different smoothing techniques, were then used for VSI. Because of the noise that remained even after smoothing, staggering and operator suppression (Algorithm 2) were used.

**Figure 6:**
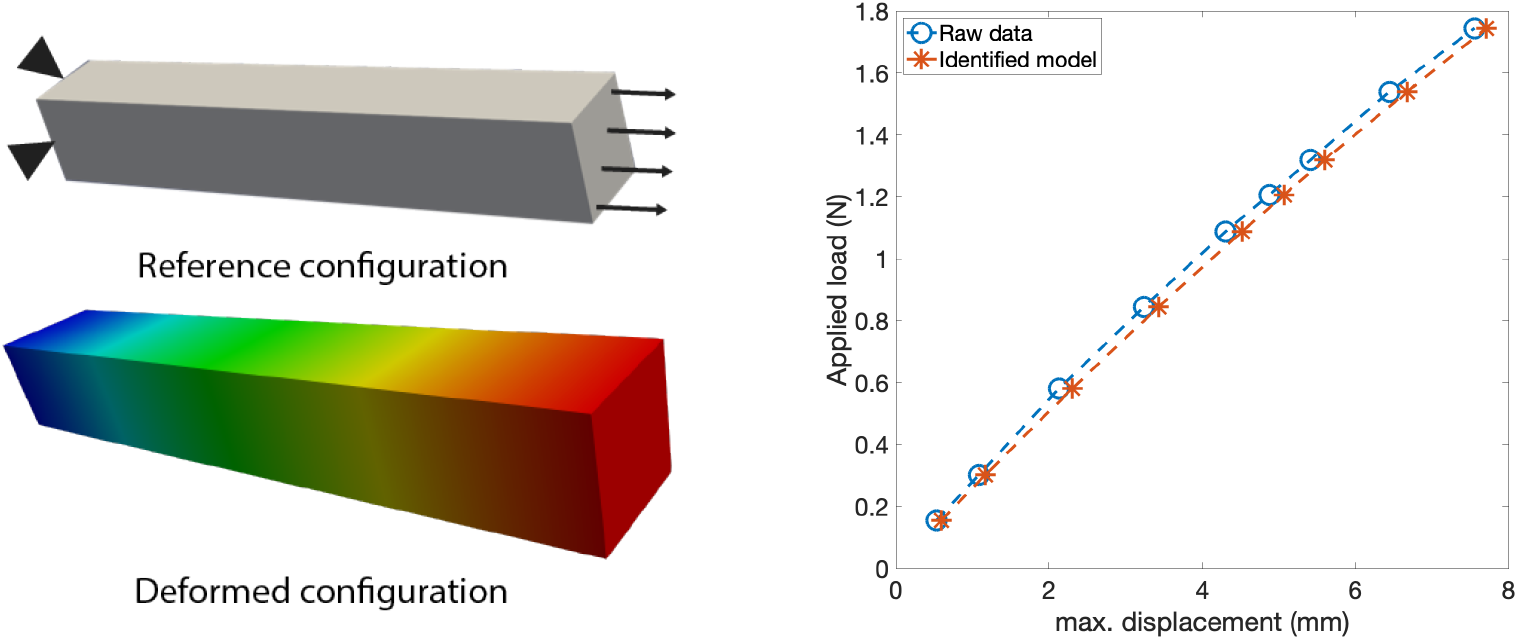
Left: reconstructed deformation from FFDC data on a soft elastomer showing the reference, and final deformed configurations. The right plot shows the load-displacement curves for the nine loading steps of the raw data (without filtering) and the results simulated by the model identified from the FFDC data (without filtering) shown in Table 5.

We first consider the strain energy density function including all the deformation mechanisms in Equation (2). For the unsmoothed FFDC data, or if smoothed by Hamming filters, the staggered VSI approach identifies the single orthotropic term of the form (*Ī*_*a*_ − 1)^2^ as the sole non-volumetric deformation mechanism. However, this manifests in either (a) a negative bulk modulus or (b) in a vanishing shear modulus, because the (*Ī*_1_ − 3), (*Ī*_2_ − 3), (*Ī*_1_ − 3)^2^ and (*Ī*_2_ − 3)^2^ terms are eliminated. These sets of inferred coefficients lead to volumetric and shear instabilities, respectively, in forward computations. For this reason we consider these conclusions of VSI to be nonphysical. These results are summarized in Table 3. However, the direct use of polynomial smoothing does not display this pathology, and identifies physically admissible mechanisms of volumetric and shear deformation. For the data subject to Hamming filters, we therefore reran Algorithms 1 and 2 with suppression of the orthotropic mechanism as described in Section 3.3.1. The corresponding results appear in Table 4. For displacement data without filtering, and with the Hamming filter of width 1^3^ on ***u***^d^, the final results remain nonphysical, with a negative bulk modulus even with operator suppression. However, we get physically meaningful results using the remaining kernels and with polynomial smoothing on ***u***^d^, as well as for all Hamming filters with ***F*** ^d^.

**Table 3:**
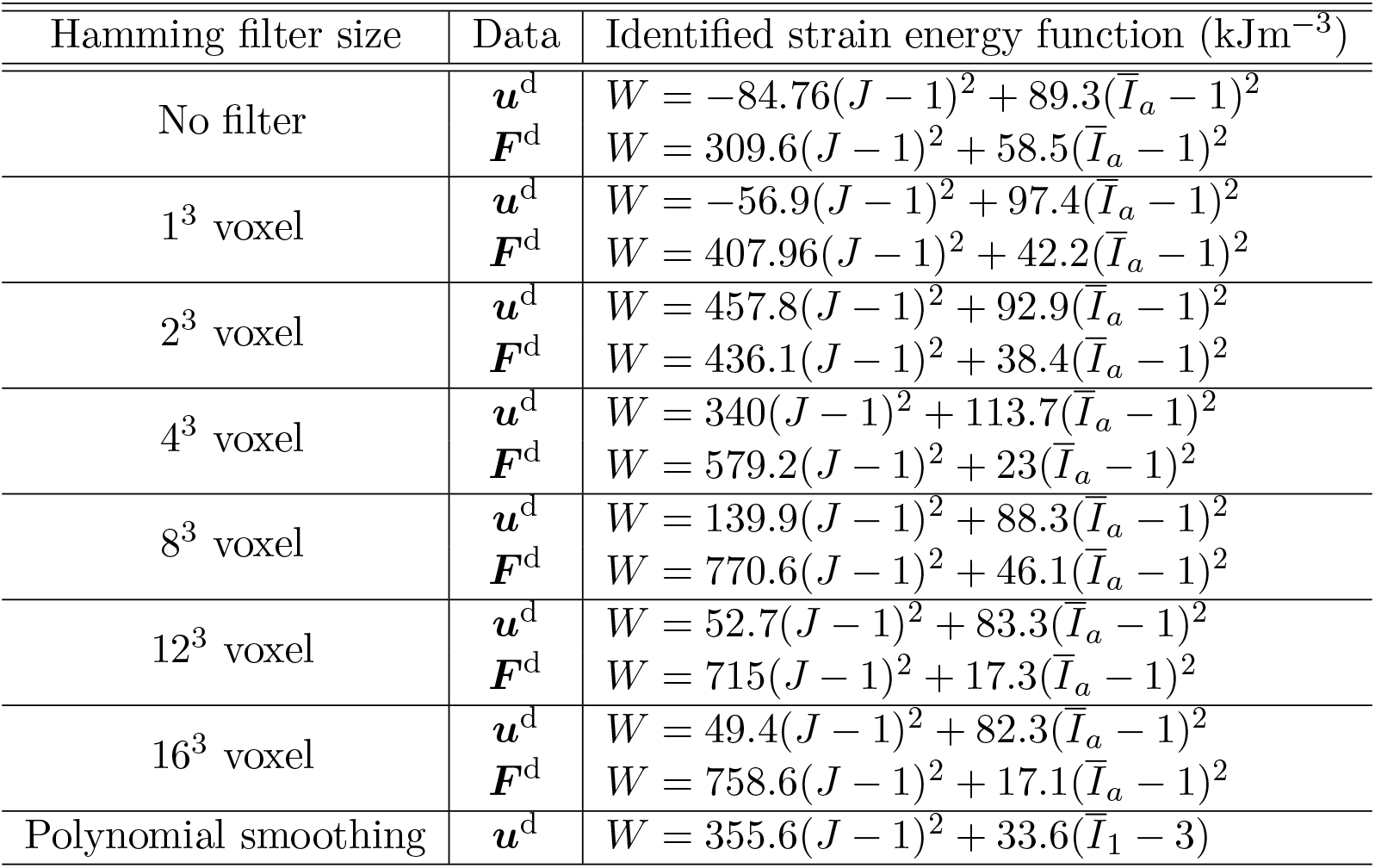
VSI results for the rectangular prism-shaped soft elastomer without suppression of any of the deformation mechanisms in Equation (2).

| Hamming filter size | Data | Identified strain energy function ($\text{kJm}^{-3}$) |
| --- | --- | --- |
| No filter | $\mathbf{u}^d$<br>$\mathbf{F}^d$ | $W = -84.76(J-1)^2 + 89.3(\bar{I}_a - 1)^2$<br>$W = 309.6(J-1)^2 + 58.5(\bar{I}_a - 1)^2$ |
| $1^3$ voxel | $\mathbf{u}^d$<br>$\mathbf{F}^d$ | $W = -56.9(J-1)^2 + 97.4(\bar{I}_a - 1)^2$<br>$W = 407.96(J-1)^2 + 42.2(\bar{I}_a - 1)^2$ |
| $2^3$ voxel | $\mathbf{u}^d$<br>$\mathbf{F}^d$ | $W = 457.8(J-1)^2 + 92.9(\bar{I}_a - 1)^2$<br>$W = 436.1(J-1)^2 + 38.4(\bar{I}_a - 1)^2$ |
| $4^3$ voxel | $\mathbf{u}^d$<br>$\mathbf{F}^d$ | $W = 340(J-1)^2 + 113.7(\bar{I}_a - 1)^2$<br>$W = 579.2(J-1)^2 + 23(\bar{I}_a - 1)^2$ |
| $8^3$ voxel | $\mathbf{u}^d$<br>$\mathbf{F}^d$ | $W = 139.9(J-1)^2 + 88.3(\bar{I}_a - 1)^2$<br>$W = 770.6(J-1)^2 + 46.1(\bar{I}_a - 1)^2$ |
| $12^3$ voxel | $\mathbf{u}^d$<br>$\mathbf{F}^d$ | $W = 52.7(J-1)^2 + 83.3(\bar{I}_a - 1)^2$<br>$W = 715(J-1)^2 + 17.3(\bar{I}_a - 1)^2$ |
| $16^3$ voxel | $\mathbf{u}^d$<br>$\mathbf{F}^d$ | $W = 49.4(J-1)^2 + 82.3(\bar{I}_a - 1)^2$<br>$W = 758.6(J-1)^2 + 17.1(\bar{I}_a - 1)^2$ |
| Polynomial smoothing | $\mathbf{u}^d$ | $W = 355.6(J-1)^2 + 33.6(\bar{I}_1 - 3)$ |

**Table 4:**
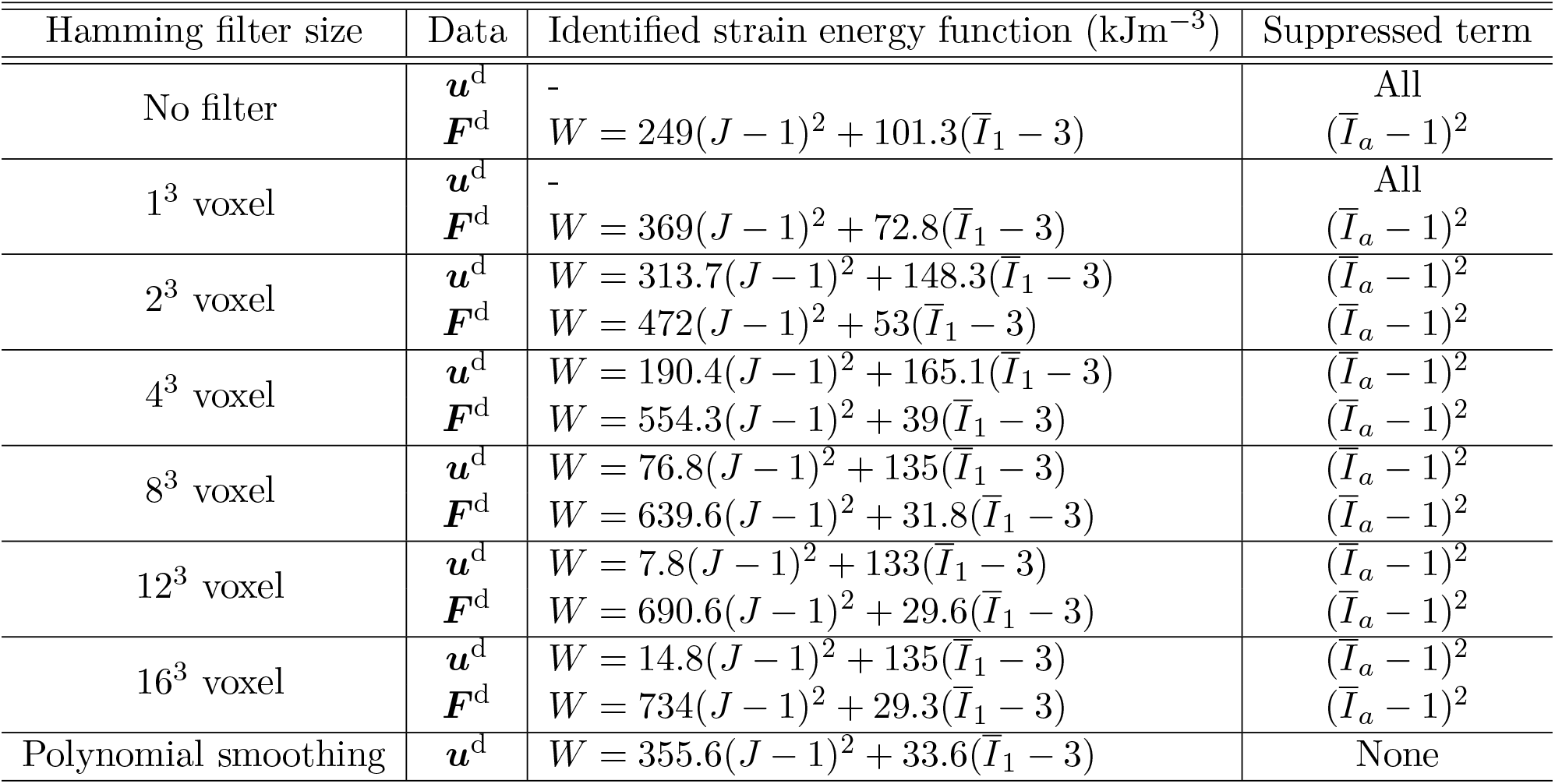
VSI results for the rectangular prism-shaped soft elastomer with operator suppression. The identified strain energy density function has volumetric and isochoric terms of the form 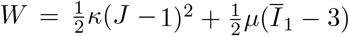

| Hamming filter size | Data | Identified strain energy function ($\text{kJm}^{-3}$) | Suppressed term |
| --- | --- | --- | --- |
| No filter | $\mathbf{u}^d$<br>$\mathbf{F}^d$ | -<br>$W = 249(J-1)^2 + 101.3(\bar{I}_1 - 3)$ | All<br>$(\bar{I}_a - 1)^2$ |
| $1^3$ voxel | $\mathbf{u}^d$<br>$\mathbf{F}^d$ | -<br>$W = 369(J-1)^2 + 72.8(\bar{I}_1 - 3)$ | All<br>$(\bar{I}_a - 1)^2$ |
| $2^3$ voxel | $\mathbf{u}^d$<br>$\mathbf{F}^d$ | $W = 313.7(J-1)^2 + 148.3(\bar{I}_1 - 3)$<br>$W = 472(J-1)^2 + 53(\bar{I}_1 - 3)$ | $(\bar{I}_a - 1)^2$<br>$(\bar{I}_a - 1)^2$ |
| $4^3$ voxel | $\mathbf{u}^d$<br>$\mathbf{F}^d$ | $W = 190.4(J-1)^2 + 165.1(\bar{I}_1 - 3)$<br>$W = 554.3(J-1)^2 + 39(\bar{I}_1 - 3)$ | $(\bar{I}_a - 1)^2$<br>$(\bar{I}_a - 1)^2$ |
| $8^3$ voxel | $\mathbf{u}^d$<br>$\mathbf{F}^d$ | $W = 76.8(J-1)^2 + 135(\bar{I}_1 - 3)$<br>$W = 639.6(J-1)^2 + 31.8(\bar{I}_1 - 3)$ | $(\bar{I}_a - 1)^2$<br>$(\bar{I}_a - 1)^2$ |
| $12^3$ voxel | $\mathbf{u}^d$<br>$\mathbf{F}^d$ | $W = 7.8(J-1)^2 + 133(\bar{I}_1 - 3)$<br>$W = 690.6(J-1)^2 + 29.6(\bar{I}_1 - 3)$ | $(\bar{I}_a - 1)^2$<br>$(\bar{I}_a - 1)^2$ |
| $16^3$ voxel | $\mathbf{u}^d$<br>$\mathbf{F}^d$ | $W = 14.8(J-1)^2 + 135(\bar{I}_1 - 3)$<br>$W = 734(J-1)^2 + 29.3(\bar{I}_1 - 3)$ | $(\bar{I}_a - 1)^2$<br>$(\bar{I}_a - 1)^2$ |
| Polynomial smoothing | $\mathbf{u}^d$ | $W = 355.6(J-1)^2 + 33.6(\bar{I}_1 - 3)$ | None |

**Table 5:** Parameters corresponding to isotropic response with bulk and shear moduli *κ* and *µ* that are further refined by PDE-constrained minimization. The initial guesses are the results of VSI using the data with the corresponding filter on ***u***^d^, or with second-order polynomial smoothing. The errors are reported for the ninth load step and should be compared with the maximum displacement of 7.57 mm. The details of the volume-averaged *L*^2^-error for each load step are summarized in Table 9 in the Appendix.

| Hamming filter size | $\kappa$ (kPa) | $\mu$ (kPa) | Vol. ave. $L_2$-error(mm) |
| --- | --- | --- | --- |
| No filter | 4064 | 60.4 | <0.11 |
| $1^3$ voxel | 3846 | 60.4 | <0.1 |
| $2^3$ voxel | 3906 | 60.4 | <0.1 |
| $4^3$ voxel | 3891.2 | 60.6 | <0.1 |
| $8^3$ voxel | 3348.4 | 60.8 | <0.1 |
| $12^3$ voxel | 2563.6 | 61.2 | <0.1 |
| $16^3$ voxel | 85.6 | 82.2 | <0.1 |
| Polynomial smoothing | 3790 | 60.3 | <0.1 |

Following inference of the deformation mechanisms by VSI, the coefficient values, specifically the bulk and shear moduli *κ* and *µ*, were further refined by PDE-constrained minimization with adjoints as described in Section 3.4. This leads to a lower shear modulus and a notably higher bulk modulus–see Table 5. VSI combined with PDE-constrained minimization thus confirms that the material is indeed nearly incompressible. The load-displacement curve of the inferred model matched the data very well as shown in the right plot in Figure 6. For the values of *κ* and *µ* corresponding to the Hamming filters with kernel size ≤ 4^3^ voxels, and the second-order polynomial smoothing in Table 5, the Poisson ratios range from *ν* = 0.492 to *ν* = 0.4926 when calculated in the infinitesimal strain regime. However, with wider Hamming filters, a more compressible response is inferred; the Poisson ratios decline to 0.491 and 0.4881 for the 8^3^ and 12^3^ Hamming filters. For comparison, results with VFM [14] applied to these FFDC data yielded *µ* ∼ 55.6 − 56.6 kPa. We then computed the volume-averaged *L*^2^-error defined as

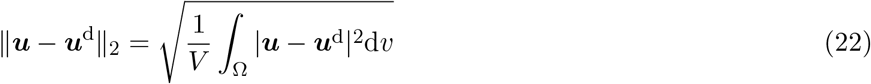

for each load step. As the applied loads increase, higher maximum displacements are obtained in the FFDC data. The corresponding volume-averaged *L*^2^-error after VSI and PDE-constrained optimization also grows with applied load; however its ratio to the maximum displacement decreases. This is because the PDE-constrained minimization places greater weightage on the load steps with higher displacement, and therefore higher error, as it minimizes the overall loss. Furthermore, focusing on the final load step, we note that while the volume-averaged *L*^2^-error is ≲ 1% for Hamming filters with kernel size ≤4^3^ voxels, and for quadratic polynomial smoothing, it increases for wider Hamming filters. The volume-averaged *L*^2^-errors corresponding to each load are summarized in Table 9 in the Appendix.

In order to arrive at confidence intervals on the inferred model parameters, we ran 300 steps of Markov Chain Monte Carlo (MCMC) sampling for varying *κ* and *µ* on a Cartesian grid in the neighborhood of their values reported in Table 5 for each data set with filtering or polynomial smoothing. For all data sets except the case with Hamming filter 16^3^, we uniformly sampled *κ* ∈ [1000, 6000] kPa and *µ* ∈ [60, 66] kPa. For data with the Hamming filter of kernel size 16^3^ voxels, we sampled *κ* ∈ [60, 150] and *µ* ∈ [60, 66]. These ranges were chosen to encompass suitable sampling intervals containing the corresponding values reported in Table 5. Figure 7, shows contours with the indicated ratios of volume-averaged *L*^2^-error with respect to the minimum error attained on the innermost contour. As seen, the confidence intervals span a wider range of *κ* than of *µ*. Furthermore, while the confidence intervals are about the same for data sets with Hamming filters of kernel size ≤ 4^3^ voxels, they increase for larger Hamming filters and for quadratic polynomial smoothing. Also note that the upper bound of the confidence intervals for *κ* exceeds the sample range for Hamming filters with kernel size of 16^3^ voxels.

**Figure 7:**
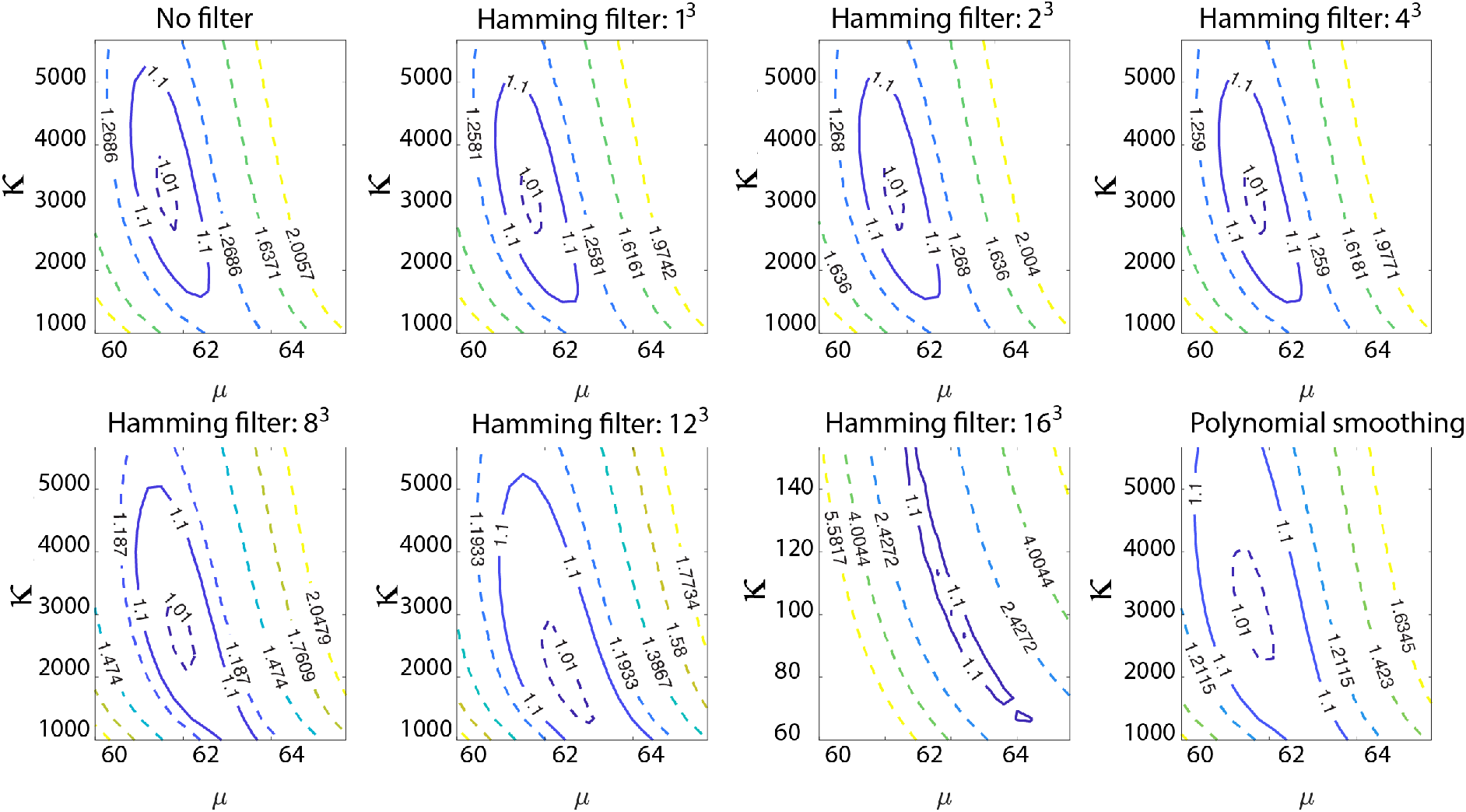
Contours of the ratio of volume averaged *L*^2^-error with sampling over *κ* and *µ* for the rectangular specimen. The ratio of error corresponding to each contour with respect to the error for the innermost contour is reported. This ratio for the solid curves is 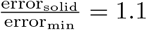.

### 4.3 FFDC data on an S-shaped soft elastomer

As the final study of inference in this communication, we applied VSI to FFDC data obtained for an “S-shaped” soft elastomer as a surrogate for irregularly shaped soft biological tissue. For this experiment, we created a custom S-shaped laser-printed mold of constant cross-section and similarly cast a solid silicone shape, but with a higher concentration of platinum, the active cross-linking ingredient (20A formulation; Dragon Skin, Smooth-On Inc., Macungie, PA). An increased cross-linking agent concentration results in a higher resulting cross-link density, which in turn increases the material stiffness; hence, we expect our shear modulus estimates to be higher.

We again fixed the displacement on one face, and stretched the opposite face longitudinally in six increments by specifying the maximum displacement to be 0.99, 1.48, 1.99, 3.02, 4.02 and 5.06 mm. The resultant forces were: 1.06, 1.58, 2.06, 2.99, 3.87 and 4.70 N, respectively, and were distributed with close to uniform traction as the elastomer was only slightly “S-shaped”. The details of the experimental configuration can be found in Ref. [14]. The reference configuration, boundary conditions and reconstructed deformed configuration for the final load increment (of six) appear in Figure 8. The displacement vector field and the deformation gradient tensor field were directly obtained from FFDC, and Hamming filters were applied with different kernel sizes. Polynomial smoothing was also carried out on the displacement data, and were best fit by globally quintic forms. As previously, the operators in Equation (3) corresponding to deformation mechanisms in Equation (2) were constructed with the traction term as the label in Equation (9).

**Figure 8:**
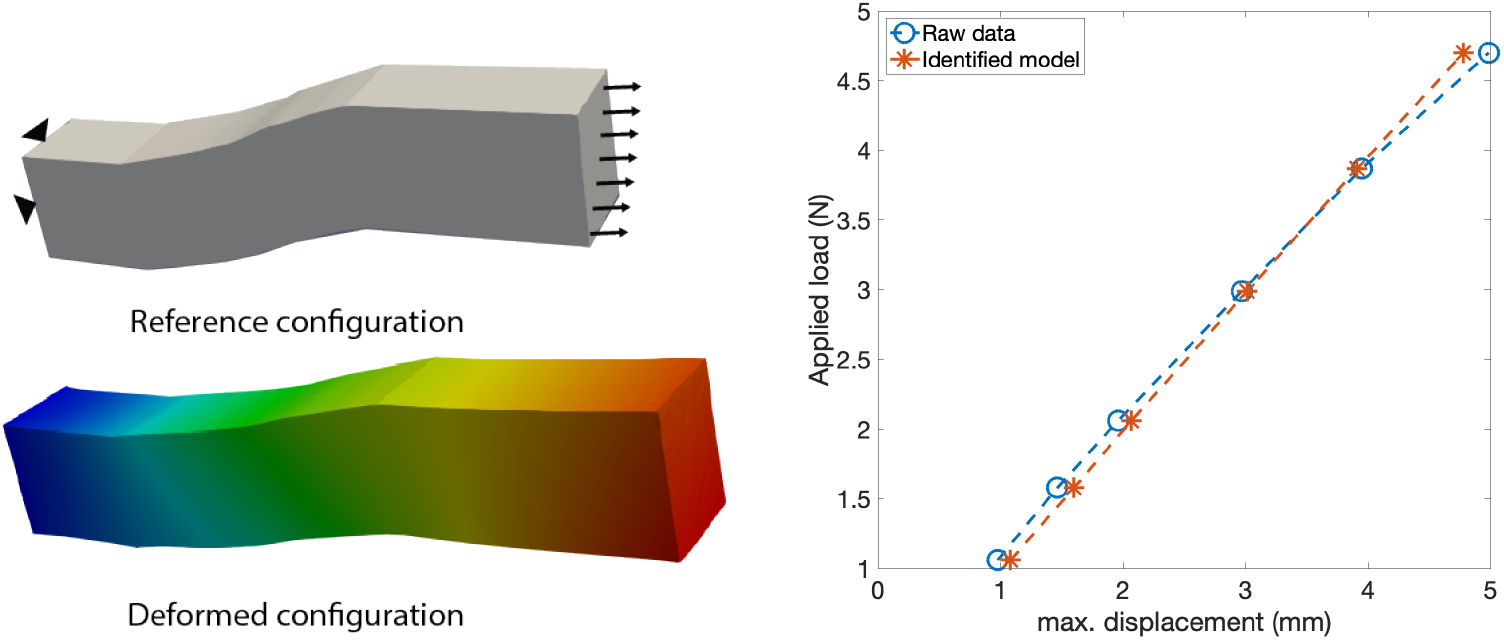
Reconstructed deformation from data on an S-shaped soft elastomer showing the reference, and final deformed configurations. The right plot shows the load-displacement curves for the six loading steps of the raw data (without filtering) and the results simulated by the model identified from the raw data (without filtering) shown in Table 8.

Following the approach used for the rectangular prismatic shape, we first included all the deformation mechanisms in Equation (2). In all cases the staggered VSI approach identified the single orthotropic term of the form (*Ī*_*a*_ − 1)^2^ as the sole non-volumetric deformation mechanism. However, this manifested in nonphysical results via either (a) a negative bulk modulus or (b) in a vanishing shear modulus, because the (*Ī*_1_ − 3), (*Ī*_2_ − 3), (*Ī*_1_ − 3)^2^ and (*Ī*_2_ − 3)^2^ terms were eliminated. Both situations imply a mechanically unstable material. These results are summarized in Table 6. We then applied Algorithms 1 and 2 with staggering and operator suppression of this orthotropic term based on linear dependence, as in Section 3.3.1. Use of unsmoothed displacement data or with the Hamming filter kernel size of 1^3^ voxels on ***u***^d^ still led to nonphysical final results with a negative bulk modulus even when operator suppression was used. But the use of field data on the deformation gradient tensor with the Hamming filter and quintic polynomial smoothing on the displacement data led to physically meaningful results (See Table 7).

**Table 6:**
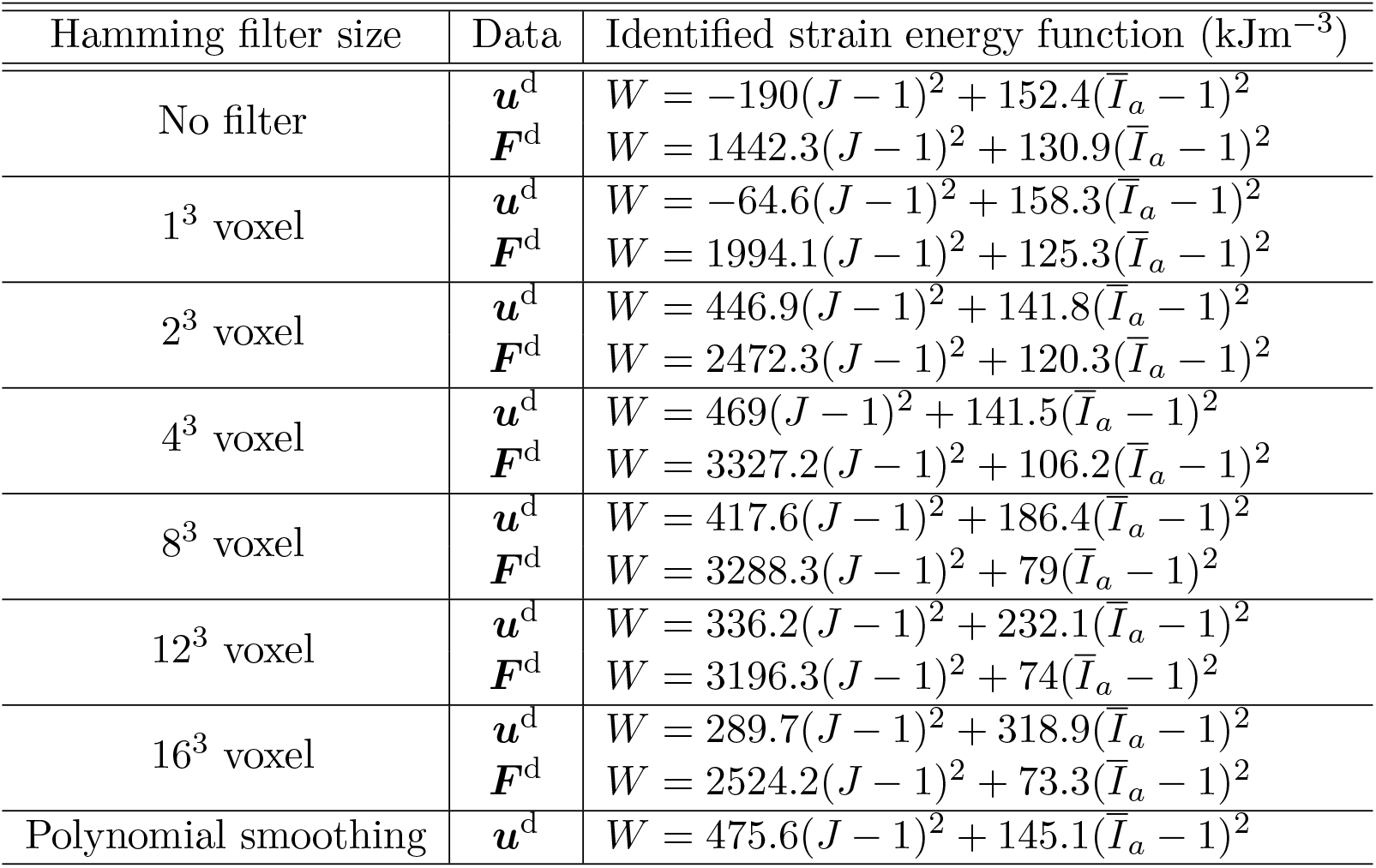
VSI results for the S-shaped soft elastomer without suppression of any of the deformation mechanisms in (2).

**Table 7:**
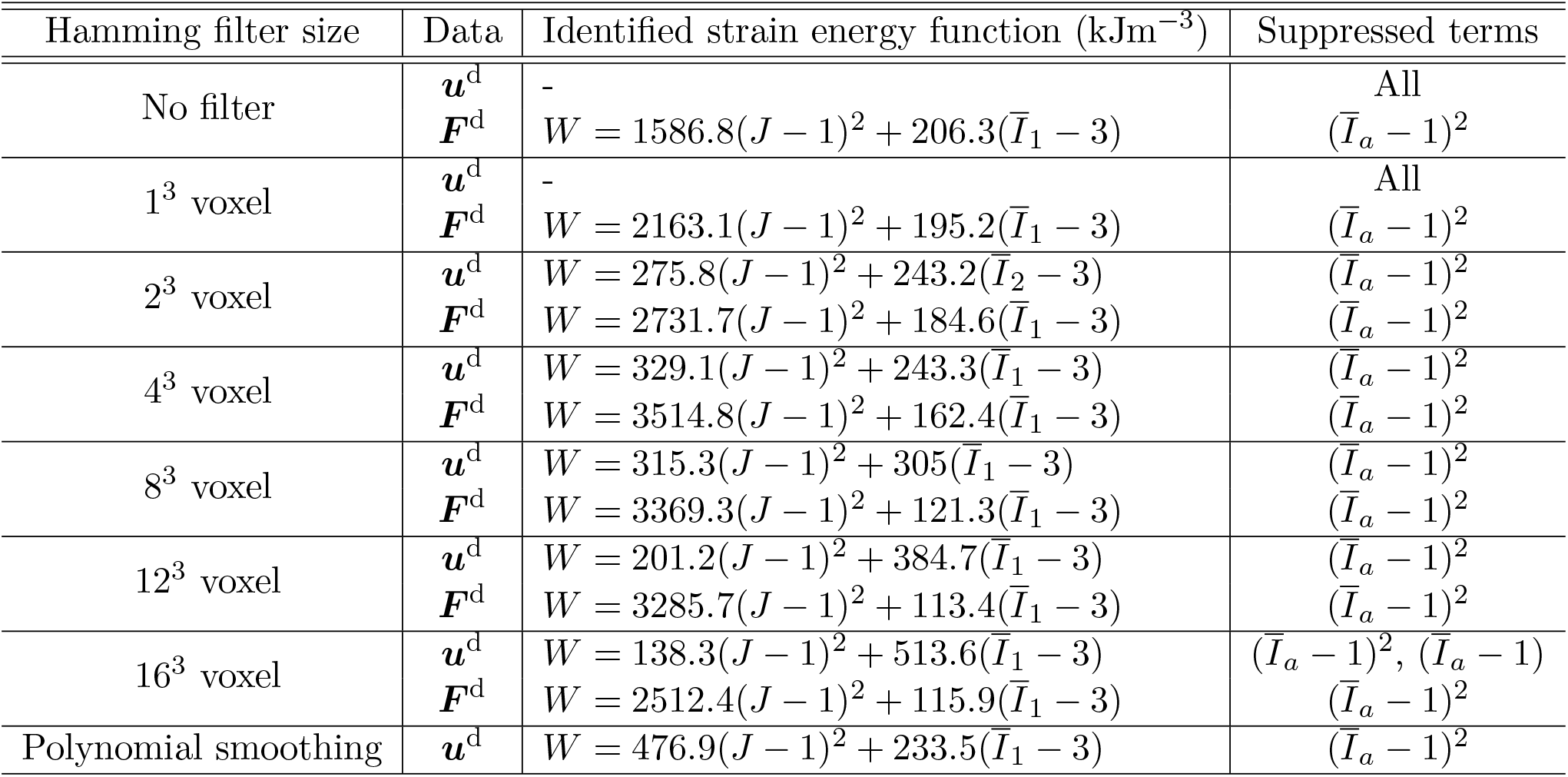
VSI results for the S-shaped soft elastomer with operator suppression. The identified strain energy density function has volumetric and isochoric terms of the form 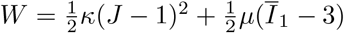

| Hamming filter size | Data | Identified strain energy function (kJm<sup>-3</sup>) | Suppressed terms |
| --- | --- | --- | --- |
| No filter | $\mathbf{u}^d$ | - | All |
| | $\mathbf{F}^d$ | $W = 1586.8(J - 1)^2 + 206.3(\bar{I}_1 - 3)$ | $(\bar{I}_a - 1)^2$ |
| 1<sup>3</sup> voxel | $\mathbf{u}^d$ | - | All |
| | $\mathbf{F}^d$ | $W = 2163.1(J - 1)^2 + 195.2(\bar{I}_1 - 3)$ | $(\bar{I}_a - 1)^2$ |
| 2<sup>3</sup> voxel | $\mathbf{u}^d$ | $W = 275.8(J - 1)^2 + 243.2(\bar{I}_2 - 3)$ | $(\bar{I}_a - 1)^2$ |
| | $\mathbf{F}^d$ | $W = 2731.7(J - 1)^2 + 184.6(\bar{I}_1 - 3)$ | $(\bar{I}_a - 1)^2$ |
| 4<sup>3</sup> voxel | $\mathbf{u}^d$ | $W = 329.1(J - 1)^2 + 243.3(\bar{I}_1 - 3)$ | $(\bar{I}_a - 1)^2$ |
| | $\mathbf{F}^d$ | $W = 3514.8(J - 1)^2 + 162.4(\bar{I}_1 - 3)$ | $(\bar{I}_a - 1)^2$ |
| 8<sup>3</sup> voxel | $\mathbf{u}^d$ | $W = 315.3(J - 1)^2 + 305(\bar{I}_1 - 3)$ | $(\bar{I}_a - 1)^2$ |
| | $\mathbf{F}^d$ | $W = 3369.3(J - 1)^2 + 121.3(\bar{I}_1 - 3)$ | $(\bar{I}_a - 1)^2$ |
| 12<sup>3</sup> voxel | $\mathbf{u}^d$ | $W = 201.2(J - 1)^2 + 384.7(\bar{I}_1 - 3)$ | $(\bar{I}_a - 1)^2$ |
| | $\mathbf{F}^d$ | $W = 3285.7(J - 1)^2 + 113.4(\bar{I}_1 - 3)$ | $(\bar{I}_a - 1)^2$ |
| 16<sup>3</sup> voxel | $\mathbf{u}^d$ | $W = 138.3(J - 1)^2 + 513.6(\bar{I}_1 - 3)$ | $(\bar{I}_a - 1)^2, (\bar{I}_a - 1)$ |
| | $\mathbf{F}^d$ | $W = 2512.4(J - 1)^2 + 115.9(\bar{I}_1 - 3)$ | $(\bar{I}_a - 1)^2$ |
| Polynomial smoothing | $\mathbf{u}^d$ | $W = 476.9(J - 1)^2 + 233.5(\bar{I}_1 - 3)$ | $(\bar{I}_a - 1)^2$ |

**Table 8:**
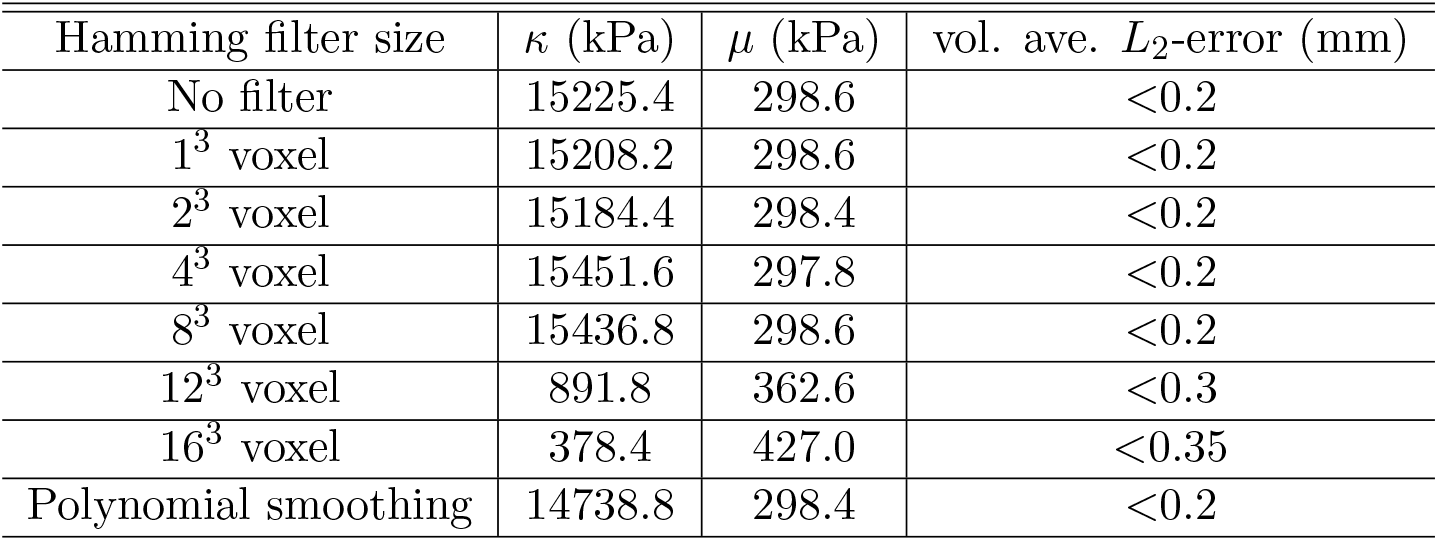
Bulk and shear moduli for the S-shaped elastomer further refined by PDE-constrained minimization with initial guesses taken from VSI results with the corresponding filter on ***u***^d^, or with polynomial smoothing. The errors are reported for the sixth load step and should be compared with the maximum displacement of 5.06 mm. The details of the volume-averaged *L*^2^-error for each load step are summarized in Table 10 in the Appendix.

Following inference of deformation mechanisms by staggered VSI with operator suppression, the identified bulk and shear moduli *κ* and *µ* were further refined by PDE-constrained minimization with adjoints. This led to a lower shear modulus and a notably higher bulk modulus with the Poisson ratio *ν*∼ 0.49 in the infinitesimal strain regime–see Table 8. For comparison, results with VFM [14] applied to these FFDC data yielded *µ* ∼ 267 kPa. Again, the load-displacement curve of the inferred model matched the data very well, with both plots displaying a slight softening response as shown on the right in Figure 8. The volume-averaged *L*^2^-error at the maximum applied load was nearly identical and ≲ 4% of the maximum displacement across data sets that were subject to Hamming filters with kernel size ≤8^3^ voxels, and for quintic polynomial smoothing, suggesting acceptability of these different imaging filter sizes for data processing purposes. In contrast, the volume-averaged *L*^2^-error increased with larger filter sizes, suggesting that overly aggressive data smoothing reduces displacement field agreement. The details of the volume-averaged *L*^2^-errors corresponding to each load are summarized in Table 10 in the Appendix. VSI combined with PDE-constrained minimization thus confirms that the material is indeed nearly incompressible and delivers consistent results for Hamming filtered data on ***F*** ^d^, and polynomial smoothing of ***u***^d^.

**Table 9:** Detailed results on forward predictions after VSI and PDE-constrained optimization for the rectangular elastomer.

| Raw data |  |  |  |  |  |  |  |  |  |
| --- | --- | --- | --- | --- | --- | --- | --- | --- | --- |
| Applied load (N) | 0.16 | 0.30 | 0.58 | 0.84 | 1.09 | 1.21 | 1.32 | 1.54 | 1.74 |
| max. displacements(mm) | 0.55 | 1.09 | 2.14 | 3.25 | 4.31 | 4.89 | 5.43 | 6.46 | 7.57 |
| Identified model using data with different filtering |  |  |  |  |  |  |  |  |  |
| No filter |  |  |  |  |  |  |  |  |  |
| Vol. ave. $L_2$-error(mm) | 0.042 | 0.057 | 0.097 | 0.1 | 0.11 | 0.09 | 0.086 | 0.09 | 0.073 |
| ratio to max. disp. | 7.64% | 5.23% | 4.53% | 3.08% | 2.55% | 1.84% | 1.58% | 1.39% | 0.96% |
| Hamming filter: $1^3$ voxel | | | | | | | | | |
| Vol. ave. $L_2$-error(mm) | 0.04 | 0.06 | 0.1 | 0.1 | 0.1 | 0.1 | 0.9 | 0.1 | 0.076 |
| ratio to max. disp. | 7.27% | 5.5% | 4.67% | 3.08% | 2.32 % | 2.04% | 1.66% | 1.55% | 1.% |
| Hamming filter: $2^3$ voxel | | | | | | | | | |
| Vol. ave. $L_2$-error(mm) | 0.04 | 0.06 | 0.095 | 0.1 | 0.1 | 0.095 | 0.09 | 0.095 | 0.074 |
| ratio to max. disp. | 7.27% | 5.5% | 4.44% | 3.08% | 2.32 % | 1.94% | 1.66% | 1.47% | 0.98% |
| Hamming filter: $4^3$ voxel | | | | | | | | | |
| Vol. ave. $L_2$-error(mm) | 0.04 | 0.05 | 0.09 | 0.096 | 0.1 | 0.09 | 0.084 | 0.09 | 0.076 |
| ratio to max. disp. | 7.27% | 4.59% | 4.21% | 2.95% | 2.32 % | 1.84% | 1.55% | 1.39% | 1.% |
| Hamming filter: $8^3$ voxel | | | | | | | | | |
| Vol. ave. $L_2$-error(mm) | 0.04 | 0.05 | 0.1 | 0.1 | 0.1 | 0.1 | 0.1 | 0.1 | 0.1 |
| ratio to max. disp. | 7.27% | 4.59% | 4.67% | 3.08% | 2.32 % | 2.04% | 1.84% | 1.55% | 1.32% |
| Hamming filter: $12^3$ voxel | | | | | | | | | |
| Vol. ave. $L_2$-error(mm) | 0.04 | 0.05 | 0.1 | 0.1 | 0.13 | 0.12 | 0.13 | 0.14 | 0.15 |
| ratio to max. disp. | 7.27% | 4.59% | 4.67% | 3.08% | 3.02 % | 2.45% | 2.39% | 2.17% | 1.98% |
| Hamming filter: $16^3$ voxel | | | | | | | | | |
| Vol. ave. $L_2$-error(mm) | 0.04 | 0.06 | 0.1 | 0.1 | 0.1 | 0.1 | 0.13 | 0.14 | 0.15 |
| ratio to max. disp. | 7.27% | 5.5% | 4.67% | 3.08% | 2.32 % | 2.04% | 2.39% | 2.17% | 1.98% |
| Second-order polynomial smoothing |  |  |  |  |  |  |  |  |  |
| Vol. ave. $L_2$-error(mm) | 0.03 | 0.055 | 0.9 | 0.096 | 0.1 | 0.08 | 0.08 | 0.08 | 0.067 |
| ratio to max. disp. | 5.45% | 5.05% | 4.21% | 2.95% | 2.32 % | 1.64% | 1.47% | 1.24% | 0.89% |

**Table 10:** Detailed results on forward predictions after VSI and PDE-constrained optimization for the S-shaped elastomer

| Raw data |  |  |  |  |  |  |
| --- | --- | --- | --- | --- | --- | --- |
| Applied load (N) | 1.06 | 1.58 | 2.06 | 2.99 | 3.87 | 4.7 |
| max. displacements(mm) | 0.99 | 1.48 | 1.99 | 3.02 | 4.02 | 5.06 |
| Identified model using data with different filtering |  |  |  |  |  |  |
| No filter |  |  |  |  |  |  |
| Vol. ave. $L_2$-error(mm) | 0.06 | 0.1 | 0.1 | 0.16 | 0.19 | 0.2 |
| ratio to max. disp. | 6.06% | 6.76% | 5.03% | 5.3% | 4.73% | 3.95% |
| Hamming filter: $1^3$ voxel | | | | | | |
| Vol. ave. $L_2$-error(mm) | 0.06 | 0.1 | 0.1 | 0.16 | 0.19 | 0.2 |
| ratio to max. disp. | 6.06% | 6.76% | 5.03% | 5.3% | 4.73% | 3.95% |
| Hamming filter: $2^3$ voxel | | | | | | |
| Vol. ave. $L_2$-error(mm) | 0.06 | 0.1 | 0.1 | 0.16 | 0.19 | 0.2 |
| ratio to max. disp. | 6.06% | 6.08% | 5.03% | 5.3% | 4.73% | 3.95% |
| Hamming filter: $4^3$ voxel | | | | | | |
| Vol. ave. $L_2$-error(mm) | 0.06 | 0.1 | 0.1 | 0.16 | 0.19 | 0.2 |
| ratio to max. disp. | 6.06% | 6.08% | 5.03% | 5.3% | 4.73% | 3.95% |
| Hamming filter: $8^3$ voxel | | | | | | |
| Vol. ave. $L_2$-error(mm) | 0.06 | 0.1 | 0.1 | 0.16 | 0.19 | 0.2 |
| ratio to max. disp. | 6.06% | 6.08% | 5.03% | 5.3% | 4.73% | 3.95% |
| Hamming filter: $12^3$ voxel | | | | | | |
| Vol. ave. $L_2$-error(mm) | 0.07 | 0.1 | 0.12 | 0.17 | 0.2 | 0.23 |
| ratio to max. disp. | 7.07% | 6.73% | 6.03% | 5.56% | 4.93% | 4.55% |
| Hamming filter: $16^3$ voxel | | | | | | |
| Vol. ave. $L_2$-error(mm) | 0.058 | 0.1 | 0.19 | 0.26 | 0.3 | 0.35 |
| ratio to max. disp. | 5.86 % | 6.76% | 9.55% | 8.61% | 7.46 % | 6.92% |
| Second-order polynomial smoothing |  |  |  |  |  |  |
| Vol. ave. $L_2$-error(mm) | 0.06 | 0.1 | 0.1 | 0.16 | 0.19 | 0.2 |
| ratio to max. disp. | 6.06% | 6.76% | 5.03% | 5.3% | 4.73 % | 3.95% |

We repeated the MCMC sampling techniques of Section 4.2 to generate confidence intervals. For Hamming filters with kernel size ≤ 4^3^ voxels and for polynomial smoothing, we sampled *κ* ∈ [4000, 100000] kPa and *µ* ≤]280, 338] kPa. The sampling was over *κ* ∈ [200, 18000] kPa and *µ* ∈ [280, 470] kPa for Hamming filters with kernel size 12^3^ voxels, and *κ* ∈ [40, 18000] kPa and *µ* ∈ [200, 780] kPa for Hamming filters with kernel size 16^3^ voxels. As in Section 4.2, these ranges were chosen based on the PDE-constrained optimization results reported in Table 8. The contours in Figure 9 are defined identically to Section 4.2: ratios of volume-averaged *L*^2^-error relative to minimum error of the innermost contour. The confidence intervals are wider for *κ* than *µ*. For all data sets, the upper bounds of the confidence intervals over *κ* exceed the sampling range. Since the error remains the same over each contour for arbitrarily large *κ* this suggests that these values are already detecting near-incompressibility.

**Figure 9:**
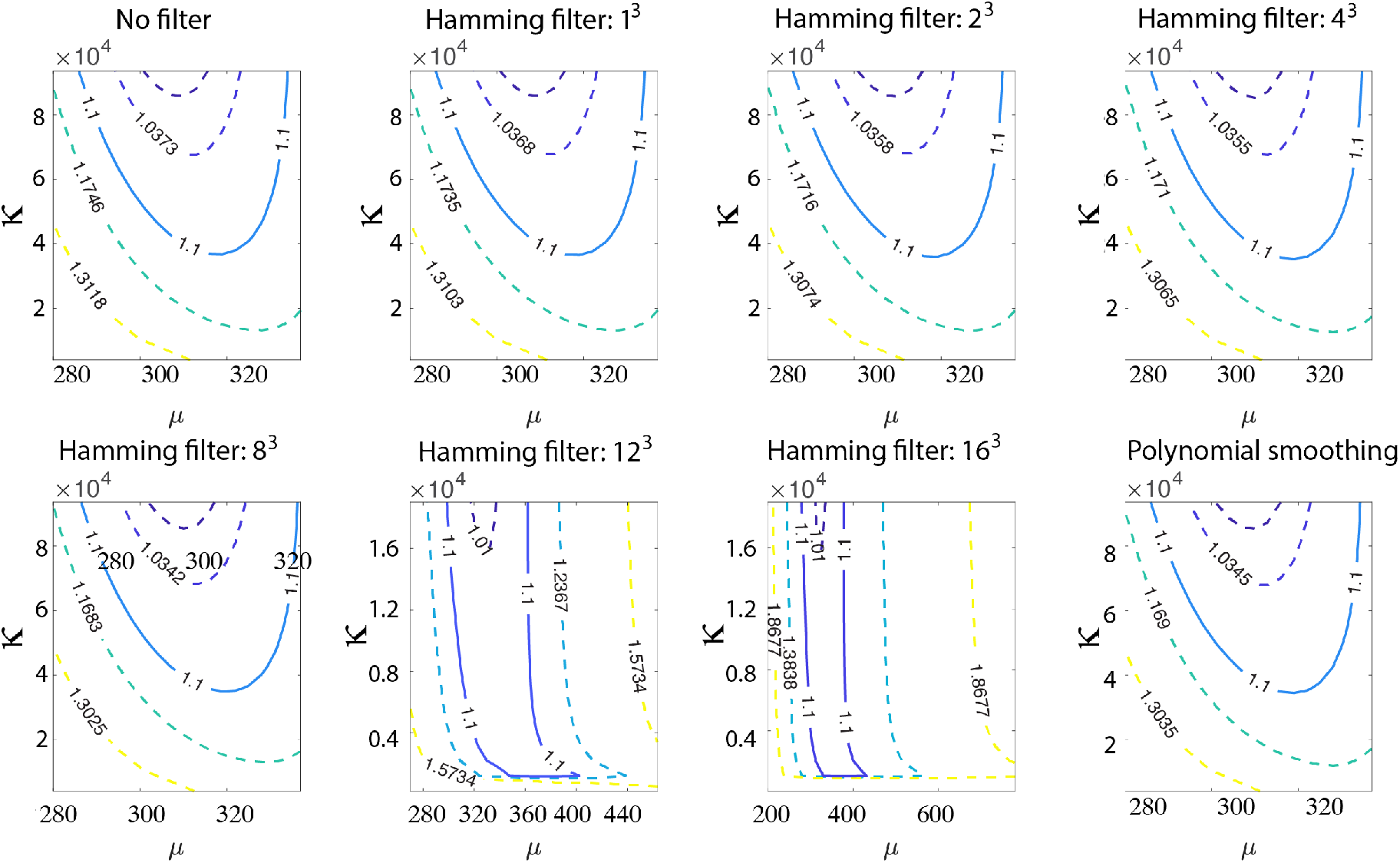
Contours of the ratio of volume averaged *L*^2^-error with sampling over *κ* and *µ* for the S-shaped specimen. The ratio of error corresponding to each contour with respect to the error for the innermost contour is reported. This ratio for the solid curves is 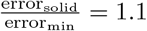.

## 5 Discussion

This work represents the first foray into solid mechanics of soft biological tissue and polymers using the framework of Variational System Identification, originally developed for inference of the physics of pattern formation in materials [37, 36, 38], and recently used to explore identification of the canonical equations of infectious disease modelling [39]. Guided by the modern treatment of soft tissue continuum mechanics at finite strain, we have based our development of VSI on hyperelasticity. We have approached constitutive modelling in the spirit of the inference of deformation mechanisms from data. Accordingly, we have considered a range of deformation mechanisms in the strain energy density function. The premise of VSI is to then use data-driven methods of system inference to find an expressive, yet efficient representation that explains the physics as constrained by the data, and in so doing, to pinpoint the deformation mechanisms that the material admits. Recognizing the depth of existing insight to the mechanics of soft material (including our own work in this area), we have restricted ourselves to volumetric, isochoric, isotropic, and orthotropic mechanisms (Equation (2)) in this, our first communication on the problem. With the hyperelastic treatment, this directly leads to algebraic operators for the first Piola-Kirchhoff stress tensor, which features in the governing equation of stress equilibrium (balance of linear momentum). This governing partial differential equation is the form in which the considered mechanisms are then sorted and culled by VSI. Unlike our earlier work on the discovery of physics in pattern forming materials, the differential operators are not in question here.

An algorithm centered on stepwise regression and the application of the statistical *F* -test is at the heart of VSI. Here we also introduced the staggered VSI approach, guided by the recognition that noise could drown out physical constraints such as the near incompressibility in the signal. Another injection of physical insight was introduced by first assuming perfect incompressibility to eliminate the volumetric term in a predictor step. This allows the robust inference of the isochoric, isotropic (shear) mechanisms, following which the reintroduction of the volumetric mechanism as a candidate led to its inference, as well, with a finite bulk modulus. The staggered VSI approach is incorporated in Algorithm 2 and leverages the core Algorithm 1 of stepwise regression and the *F* -test.

The mechanics of soft biological tissue and polymeric materials presents challenges: some encountered previously and others novel. Noise in the data, which is nearly unavoidable in experiments, is the first of these challenges as seen in the results with synthetic data on which we superposed thermal-like Gaussian noise (Table 1). As expected, the naïve application of VSI yields very accurate inference in the absence of noise but degrades to nonphysical results, indicated by the introduction of material instabilities with increasing noise. We did not scale the noise by the reported displacement in accordance with displacement-encoding MRI-based observations. This step would lead to greater discrepancies in the reported inference and may be descriptive for, or applicable to, full-field displacement methods such as image- or volume-based correlation. Here, guided by the broadly applicable principle of stability we introduced operator suppression resulting in successful inference, albeit with some loss in fidelity in the nearly incompressible response expected (Tables 1 and 2).

Taking this forward to real data on the tensile response of the rectangular prismatic elastomeric specimen as a surrogate for soft biological tissue, we used suppression of the orthotropic deformation mechanism, because it was found to survive stepwise regression but led to nonphysical results implying instability of the volumetric and shear responses. However, the combination of staggering and operator suppression via Algorithm 2 delivered physically meaningful inference of deformation mechanisms that parsimoniously identified volumetric and shear mechanisms, rejecting five other candidates of isochoric and orthotropic response. In this application, we also found Hamming filtering of the deformation gradient field and polynomial smoothing of the displacement to be effective in reducing the noise of the raw data. As a final step, PDE-constrained optimization with adjoints used for gradients resulted in highly refined final coefficient values for the bulk and shear modulus that indeed identified a nearly incompressible response. The resulting error in the forward solution computed with the inferred deformation mechanisms and parameters resulted in volume-averaged *L*^2^-error ≲ 3.5% when averaged over the nine steps of loading.

The strong suit of this work is that the experimental characterization via full field displacement capture makes available three-dimensional deformation field data for experimental characterization, and VSI identifies deformation mechanisms while maintaining efficiency of representation with these data. The rectangular prismatic specimen does indeed provide us with three-dimensional data, although the prominent deformation is along the tensile direction. The final data set, coming from an S-shaped elastomeric specimen surrogate for soft biological tissue, was loaded parallel to its longer dimension. However, this specimen yielded relatively larger and nonuniform lateral displacements, and with more noise. In this case, the same combination as above: suppression of the orthotropic deformation mechanism that otherwise leads to a nonphysical material instability in the inferred physics, with staggered VSI and finally PDE-constrained optimization using adjoints predicts nearly incompressible response with volume-averaged *L*^2^-errors of ≲ 7.5% over the six steps of loading. The S-shaped specimen provides us with data in which the three-dimensionality of the deformation is a little more pronounced than in the rectangular prismatic specimen. While the availability of such data helps inference with VSI, as demonstrated with the synthetic data obtained using a wider array of boundary conditions and loading, this trend was countered by the higher noise in the S-shaped specimen’s data. This case thus offers some insight to the practical role of noise for different datasets.

We note that filtering of the data has its limitations: Hamming filters of up to 8^3^ voxels work well for data collection with FFDC, while larger filters overly smoothen the displacement fields. This data distortion can lead to either non-physical inferred results (vanishing or negative moduli), or to the suppression of properties such as the degree of incompressibility, in this case.

Notably, the above array of techniques in VSI combined with PDE-constrained optimization using adjoints also sheds light on optimal experimental conditions. The use of larger kernels on the Hamming filters leads to higher errors and a more compressible volumetric response. This suggests approaches by which the optimal filter size and type could be chosen for optimizing the output of a given experimental setup. In order to obtain confidence bounds on the inferred coefficients, we performed MCMC sampling in the *κ* − *µ* space around the lowest error results emerging from the PDE-constrained optimization following VSI for the rectangular and “S-shaped” specimens. Using an MCMC procedure driven by error thresholds, we have reported the spread in *κ* and *µ* for each type of smoothing used.

In materials with well-understood physical mechanisms for deformation, the FFDC + VSI procedure can be harnessed to not only determine material properties, but additionally has potential to elucidate, for example, the sources and relative prevalence of experimental errors. MRI inherently reconstructs image information from sampling magnetic response in frequency space; detector error associated with specific frequency content can be isolated by procedurally testing filter parameters as included variables. In this way, individual experiments run with identical setup parameters can have systematic effects identified independently of random (thermal) noise fluctuations. Similarly, error may in general be associated with global displacement (or load level) as in digital image and volume correlation or constructed as periodic signals as in MRI detector error; incorporating specific forms of the error into our analysis allows for testing of corresponding error-generation hypotheses which in turn serve to optimize our experimental procedure.

We note that the anisotropic mechanism considered in this work involved a chosen direction of orthotropy. However, in ongoing work, we have found VSI to be effective in inferring the axes of symmetry without having to resort to elimination from among a chosen set of mechanisms with different directions. This takes VSI out of the realm of linear regression to general nonlinear optimization in Algorithms 1 and 2. Furthermore, we have shown in recent work [39] that VSI and PDE-constrained optimization using adjoints also successfully infer spatiotemporally varying mechanisms and parameters. This result also is relevant to soft biological tissues with varying axes of symmetry. When combined with the ability to infer such axes without *a priori* inclusion of a discrete set of directions, this opens the door to inferring spatially varying, and unknown axes of anisotropy that characterize soft biological materials.

## Acknowledgements

We acknowledge the support of Defense Advanced Research Projects Agency (DARPA), under Agreement No. HR0011199002, “ Artificial Intelligence guided multi-scale multi-physics framework for discovering complex emergent materials phenomena” (ZW and KG).

## Appendix

## Footnotes

1 Regression on a loss function that includes penalization on the optimization quantities to control their magnitudes.

2 In the latter approach, one data point is omitted from each round of training and used as cross-validation. For say *n* data points this therefore involves *n* rounds of training and cross-validation.

